# Templating S100A9 amyloids on Aβ fibrillar surfaces revealed by charge detection mass spectrometry, microscopy, kinetic and microfluidic analyses

**DOI:** 10.1101/2020.05.26.116400

**Authors:** Jonathan Pansieri, Igor A. Iashchishyn, Hussein Fakhouri, Lucija Ostojić, Mantas Malisauskas, Greta Musteikyte, Vytautas Smirnovas, Matthias M. Schneider, Tom Scheidt, Catherine K. Xu, Georg Meisl, Tuomas P. J. Knowles, Ehud Gazit, Rodolphe Antoine, Ludmilla A. Morozova-Roche

## Abstract

The mechanism of amyloid co-aggregation and its nucleation process are not fully understood in spite of extensive studies. Deciphering the interactions between proinflammatory S100A9 protein and Aβ_42_ peptide in Alzheimer’s disease is fundamental since inflammation plays a central role in the disease onset. Here we use innovative charge detection mass spectrometry (CDMS) together with biophysical techniques to provide mechanistic insight into the co-aggregation process and differentiate amyloid complexes at a single particle level. Combination of mass and charge distributions of amyloids together with reconstruction of the differences between them and detailed microscopy reveals that co-aggregation involves templating of S100A9 fibrils on the surface of Aβ_42_ amyloids. Kinetic analysis further corroborates that the surfaces available for the Aβ_42_ secondary nucleation are diminished due to the coating by S100A9 amyloids, while the binding of S100A9 to Aβ_42_ fibrils is validated by a microfuidic assay. We demonstrate that synergy between CDMS, microscopy, kinetic and microfluidic analyses opens new directions in interdisciplinary research.

## Introduction

In spite of the key clinical importance of amyloid formation, the mechanisms of co-aggregation of different amyloid species remain elusive. Amyloid formation is a widespread phenomenon routed in the generic property of polypeptide chains to self-assemble into cross-β-sheet containing superstructures^1–2^ and manifested in numerous amyloid diseases^3,4^ and functional amyloids.^5,6^ Comorbidity of these diseases was reported to be linked to the co-aggregation of amyloidogenic proteins.^7,8^ In Alzheimer’s disease (AD), the amyloid-neuroinflammatory cascade is manifested in coaggregation of Aβ with proinflammatory S100A9 protein, which leads to intracellular and extracellular amyloid assembly, amyloid plaque depositions and cellular toxicity.^9^ S100A9 co-aggregates with Aβ also in traumatic brain injury, which is considered as a potential precursor state for AD.^10^ The amyloid self-assembly of Aβ was well described by the involvement of secondary nucleation pathways promoted by Aβ amyloid surface.^11^ In contrast, S100A9 undergoes nucleation dependent autocatalytic amyloid growth.^12^ There is a genuine unmet need to understand the architecture and mechanism of self-assembly leading to the formation of hetero-aggregates composed of various amyloid polypeptides. Since amyloids formed by individual polypeptides are highly polymorphic,^13–15^ their co-aggregates add up to the complexity and heterogeneity of amyloid mixture. This complex problem has been addressed previously in a number of studies – the co-assembly of Aβ^40^ and Aβ_42_ was investigated by global kinetic analysis^16^ and FTIR^17^, self-sorted supramolecular nanofibrils by in situ real-time imaging^18^, co-aggregates of wild-type α-synuclein with the familial mutant variant by dual-colour scanning for intensely fluorescent targets^19^ and Aβ_42_ peptide with analogue of islet amyloid peptide by NMR.^20^ The combination of advanced techniques, including high resolution microscopy, amyloid kinetics and microfluidic analyses and state of the art CDMS as a single particle approach, were used here to resolve this problem for Aβ_42_ and S100A9 co-assemblies.

In CDMS the mass to charge (*m/z*) and charge (*z*) of an ionized molecule are measured simultaneously, enabling to determine the molecular mass directly, i.e. without resigning to *m/z* standards.^21,22^ Robustness of the technique allows the measurement of thousands of particles within reasonable time, providing the reconstruction of molecular mass distribution. Recent advances in instrumentation, in particular use of an ion trap, have significantly decreased the detection limit arising from the low charge of biological objects.^23^ This technique in the single pass mode^21,22^ was applied to reconstruct the mass distribution of individual polypeptide fibrils.^24,25^ Here we report for the first time that by advancing the method and mapping the two-dimensional frequency and difference distributions between amyloid samples, we are able to discriminate not only between the fibrils of individual polypeptides, specifically Aβ_42_ and S100A9, but also differentiate their combined complexes. These observations are reinforced further by the morphological and statistical atomic force microscopy (AFM) analysis, demonstrating that S100A9 amyloids are indeed templated on the surface of Aβ_42_ fibrils.

The reaction kinetics analysis enables to dissect the complex amyloid co-aggregation process into the multiple microscopic events, including (i) primary nucleation, i.e. spontaneous formation of nuclei acting as initial aggregation centers; (ii) elongation, i.e. growth of existing fibrils via adding monomers to their ends and (iii) secondary nucleation, involving fibril surface catalyzed formation of additional aggregation nuclei, which can significantly increase the rate of the overall process of amyloid self-assembly.^11,16,26,27^ By using kinetic analysis and immuno-gold transmission electron microscopy (TEM), it has been shown that the blocking of secondary nucleation on Aβ fibril surface can be achieved via binding of the Brichos chaperon domain.^28,29^ Transient binding events on the fibrillar surface were demonstrated also by dSTORM and AFM.^30,31^ The results from the global kinetic analysis presented here further corroborate the suppression of secondary nucleation on Aβ_42_ fibrils by S100A9 amyloid deposits. Moreover, the microfluidic binding measurements directly demonstrate the binding of S100A9 to Aβ_42_ fibrils.

## Results and discussion

Amyloids of Aβ_42_, S100A9 and joint Aβ_42_-S100A9 were formed at 30 μM concentration of each component incubated for 24 h individually or in the mixture in PBS, pH 7.4 and 42°C using 432 orbital shaking each 10 min. AFM imaging and AFM statistical analysis (Fig. 1, S1^†^ and S2^†^) demonstrate that Aβ_42_ alone self-assembles into straight fibrils with median height in the AFM cross-sections of 4.28±0.44 nm and median length of 0.41±0.13 μm, while S100A9 forms coily, much thinner and shorter fibrils with 1.8±0.21 nm median height and 0.12±0.06 μm length, respectively. In comparison, Aβ_42_-S100A9 complexes are presented as straight fibrils with 4.66±0.7 nm median height and 0.75±0.28 μm median length, respectively (Fig. 1C,E, S1^†^and S2^†^). The AFM cross-sectional height of these fibrils is characteristic of those of Aβ_42_ incubated alone, but they become significantly longer. Importantly, they are decorated on their surfaces by coily and short filaments with the same lengths and cross-sectional heights as S100A9 fibrils incubated separately (Fig. 1C, S1C-E^†^ and S2^†^). The coily filaments were observed in the same sample separately from Aβ_42_-S100A9 complexes and they were characterized by similar length and height as the filaments decorating thick Aβ_42_ carrier fibrils. This indicates that at least some S100A9 molecules were self-assembled into individual S100A9 fibrils. Interestingly, if Aβ_42_ fibril length distribution is shifted towards higher values within the Aβ_42_-S100A9 complexes, both S100A9 fibril length and height distributions remained the same irrespectively as weather they were incubated separately, attached to Aβ_42_ fibril surface or present individually in the Aβ_42_-S100A9 mixed solution (Fig. S2^†^). These results suggest that the bulk of co-aggregated complexes is represented by Aβ_42_ amyloids templating S100A9 fibrils on their autocatalytic surfaces. The templating on fibrillar surfaces rather than block polymerization is supported by the inability for these peptides to form mixed cross-β-sheet structure within the same fibril, due to a lack of complementarity between their amino acid sequences, as shown previously by FTIR.^17^ CDMS measurements were thus critical to corroborate further this hypothesis, allowing us to evaluate the full distribution of amyloids within a given sample, which is not possible with partial sampling by AFM microscopy.

**Figure 1.**
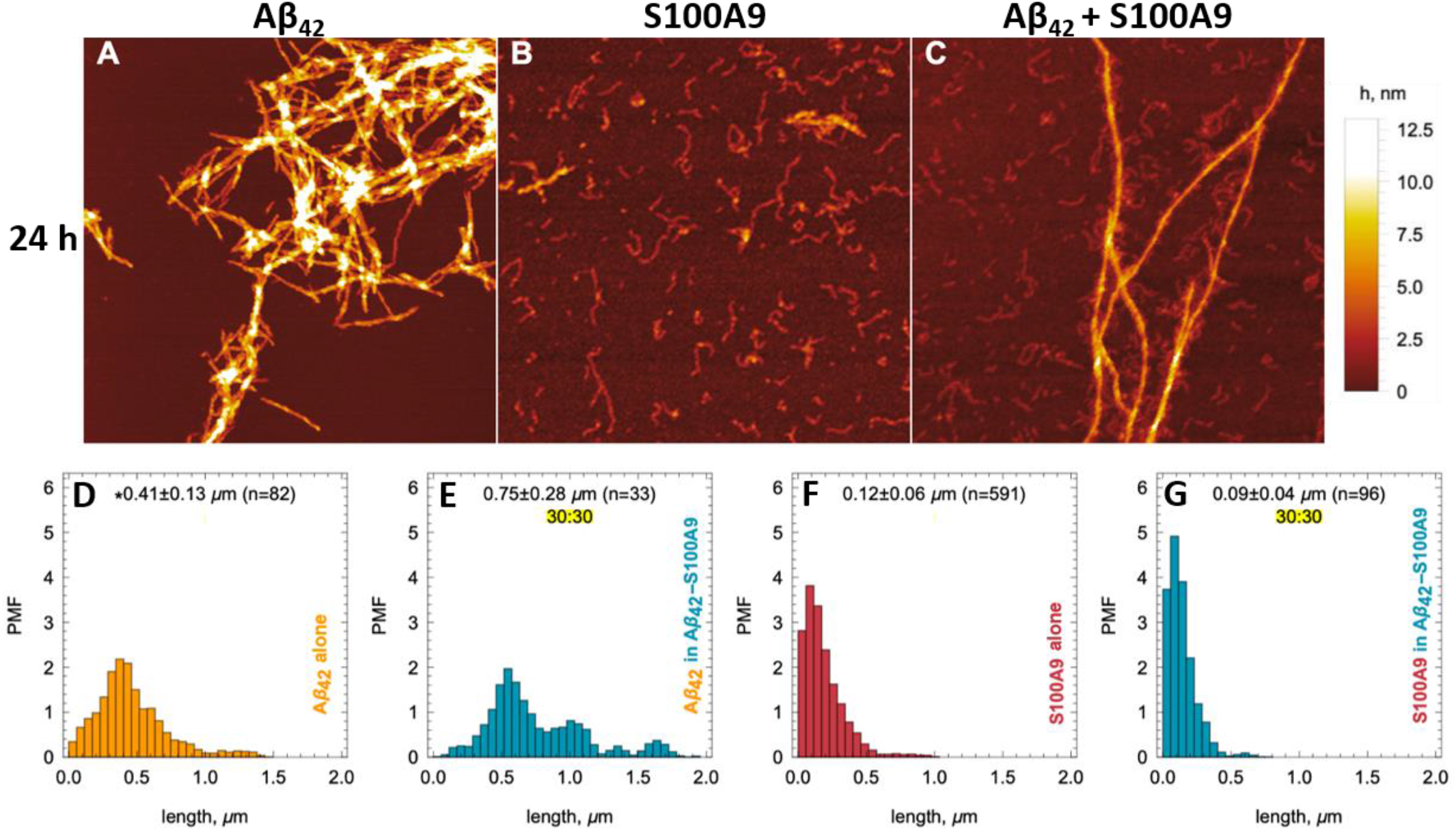
Templating S100A9 amyloids on Aβ_42_ fibrillar surface revealed by AFM. (A-C) AFM images of (A) Aβ_42_, (B) S100A9 and (C) Aβ_42_-S100A9 amyloids. Scan sizes are 2 x 2 μm. Colour scale is represented on the right. 30 μM of each polypeptide were incubated individually or in mixture with each other for 24 h in PBS, pH 7.4 and 42°C. Fibril length distributions of (D) Aβ_42_ incubated separately, (E) Aβ_42_ within Aβ_42_-S100A9 complex, (F) S100A9 incubated separately and (G) S100A9 within Aβ_42_-S100A9 complex. Probability mass function (PMF) defined as the probability of finding a fibril with a specific length is indicated along *y*-axes. Fibrillar lengths are indicated along *x*-axes. *The medians of fibril lengths with corresponding median deviations and sample sizes are shown within figures. The distributions are resampled to 10^4^ (See Material and Methods^†^).

The original CDMS data sets for the Aβ_42_, S100A9 and Aβ_42_-S100A9 samples and histograms of their molecular mass and charge distributions are shown in Fig S3^†^and 2A-C, respectively. Since the mass and charge distributions are characterized by different shapes and therefore belong to different classes of distributions, the comparison between their location metrics (mean or median values) will be biased. For example, the S100A9 fibril population is clearly represented by two sub-populations – a highly abundant low molecular mass population and an evenly distributed higher molecular mass population. Therefore, the two-dimensional frequency distributions, demonstrating the probability of finding the particle with corresponding mass and charge simultaneously and termed as frequency maps, were built up and are shown in Fig. 2E-G (described in Fig. S4^†^ and Materials and Methods^†^). These maps reveal the specific population signature for each amyloid sample and enable us to compare them with each other. The population of Aβ_42_ fibrils is characterized by proportional spread of masses and charges (Fig. 2E); the analysis of CDMS data, separately in each dimension shows that 37% Aβ_42_ fibrils fall below 80 MDa and 10% – <0.36 ke (Table S1^†^). The presence of high molecular mass particles in this distribution is likely to reflect fibril clustering, as shown in AFM and TEM images (Fig. 1A, S1A^†^, S5A^†^ and B^†^). The mass distribution for Aβ_42_ fibrils reported in this research is consistent with that reported previously.^25^

**Figure 2.**
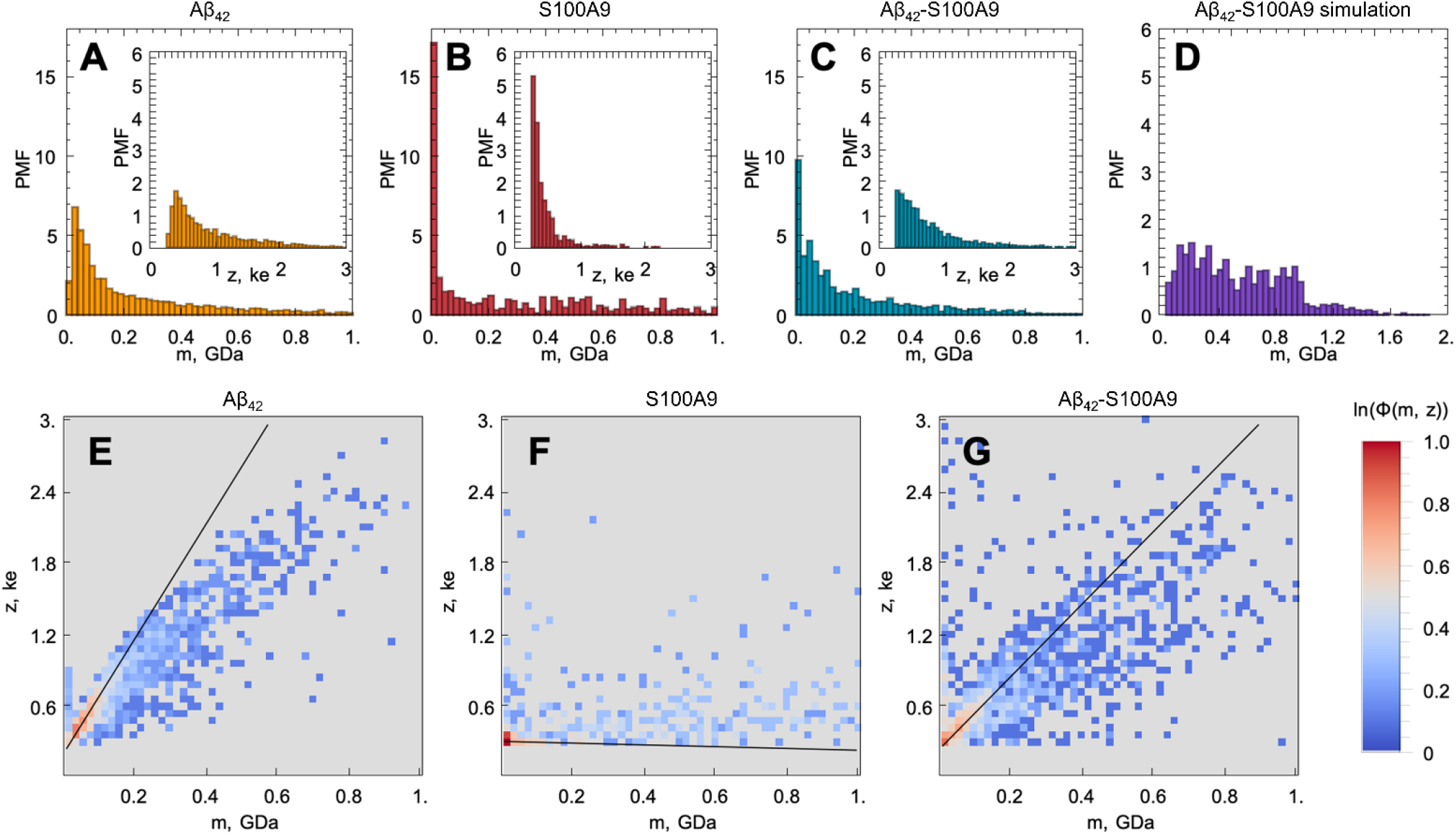
Templating S100A9 amyloids on Aβ_42_ fibrillar surface revealed by CDMS. Molecular mass (*m*) and charge (*z*, shown in insertions) distributions of (A) Aβ_42_, (B) S100A9 and (C) Aβ_42_-S100A9 amyloids. The CDMS distributions for Aβ_42_ fibrils are shown in yellow, for S100A9 fibrils – in red and for Aβ_42_-S100A9 amyloids – in blue. (D) Simulated molecular mass distribution for mixed AIϏI42 and S100A9 fibrils pre-incubated separately (shown in purple). y-axes show probability mass function (PMF) defined as the probability of finding a fibril with the corresponding m or z, which are plotted along x-axes, respectively. (E-G) CDMS populations of amyloid particles are shown as frequency maps to demonstrate the probability of finding a particle with the corresponding m and z simultaneously: (E) for Aβ_42_, (F) – S100A9 and (G) – Aβ_42_-S100A9 amyloids. Colour scale is represented on the right.

The frequency map of S100A9 fibrils (Fig. 2F) demonstrates that they are lower in masses (45% – <80 MDa), which is consistent with the morphology of these short and coily fibrils observed by AFM and TEM (Fig. 1B and S5C^†^ and D^†^), and significantly lower in charges (55% – <0.36 ke) (Table S1^†^). It is worth noting that we were able to observe such low charged population of amyloid fibrils due to the improved signal to noise ratio of the home-built CDMS instrument (Materials and Methods^†^). Indeed, the S100A9 fibrils display low charges compared to the charges on Aβ_42_ fibrils observed here and in previous experiment as well as the charges on α-synuclein and tau fibrils reported previously.^25^ The ranking of amyloid particles according to their CDMS *z*/*m* ratio for each amyloid sample (described in Materials and Methods^†^) indicates that most of the individual fibrils of Aβ_42_, S100A9 and Aβ_42_-S100A9 are lower in charge than the corresponding monomers of Aβ_42_ and S100A9, which charges were calculated using their amino acid sequence at pH 7.4 (Fig. S6^†^). This is also consistent with previous data on the shielding of monomer charges within amyloid fibrils.^24^ The population of particles with high masses in the S100A9 sample may reflect the clustering of few very flexible S100A9 fibrils into supercoils, as shown by TEM imaging (Fig. S5C^†^ and D^†^).

The frequency map of Aβ_42_-S100A9 complexes deviates from those of individual Aβ_42_ and S100A9 amyloids (Fig. 2G): 43% particles are < 80 MDa and 20% are <0.36 ke (Table S1^†^). The distribution of data points is much broader in the Aβ_42_-S100A9 frequency map and reflects partially the presence of free S100A9 fibrils in the sample as revealed by AFM (Fig. 1C, S1^†^ and S2^†^). The slope of *z* to *m* corresponding to the population of Aβ_42_-S100A9 complexes is intermediate between those for Aβ_42_ and S100A9 fibril populations, respectively, which reflects the coating of Aβ_42_ fibril surfaces by low charged S100A9 fibrils (Fig. 2E-G). The wide mass distribution may be related to the fact that S100A9 fibrils templated on Aβ_42_ amyloid surfaces make them heavier and also by blocking Aβ_42_ secondary nucleation, they promote Aβ_42_ fibril elongation, as measured by AFM (Fig. 1D and E). At the same time S100A9 coating may also make the Aβ_42_-S100A9 amyloids less prone to clumping. The presence of low *m* and high *z* complexes may reflect the population of Aβ_42_ fibrils with surface bound S100A9 monomers, since they can bind to Aβ_42_ fibrillar surface as we will discuss further.

In order to distinguish within the Aβ_42_-S100A9 sample the subpopulations of joint hetero-molecular complexes and discriminate them from the sub-populations of individual fibrillar components, such as free Aβ_42_ and S100A9 fibrils still present in this sample, we have advanced the CDMS methodology by building difference frequency distributions (described in Material and Methods^†^ and shown in Fig. S4^†^). The difference frequency distribution maps were derived by comparing the following samples: pairwise Aβ_42_-S100A9 and Aβ_42_ (Fig. S7A^†^ and D^†^); pairwise Aβ_42_-S100A9 and S100A9 (Fig. S7B^†^ and E^†^) and Aβ_42_-S100A9 *vs* pair of Aβ_42_ and S100A9 samples filtered out together (Fig. S7C^†^ and F^†^). This enables us to split the original CDMS data set into new sub-sets, demonstrating the enriched and depleted sub-populations of particles, respectively. Thus, by using this differential analysis we were able to filter out the component of interest, i.e. the sub-population of Aβ_42_-S100A9 complexes, which is clearly distinct from both Aβ_42_ and S100A9 amyloid sub-populations within the co-aggregated sample.

In addition, we have simulated the mass distributions of mixed Aβ_42_ and S100A9 fibrils formed separately and then mixed together (described in Materials and Methods^†^) and compared that with the observed CDMS mass distribution of Aβ_42_-S100A9 co-aggregates as shown in Fig. 2C and D. The simulation demonstrates that the mass distributions of co-aggregated Aβ_42_-S100A9 complexes and mixed pre-formed amyloids of Aβ_42_ and S100A9 significantly deviate from each other. While the mixed fibrils are almost evenly distributed over broad range of molecular masses, the CDMS population of Aβ_42_-S100A9 complexes displays exponential distribution. This further indicates that co-aggregation leads to a new type of joint complex formation.

In order to shed light on the co-aggregation mechanisms of Aβ_42_-S100A9 complexes, we performed the kinetics analysis of S100A9 aggregation alone and Aβ_42_ in the presence of increasing S100A9 concentrations using a thioflavin T (ThT) fluorescence assay (Fig. 3A-C and S8A^†^). Aβ_42_ fibrillation has been extensively studied previously and shown that it is governed not only by primary nucleation, but also by the secondary nucleation on the surface of already formed fibrils.^11,32^ By contrast, S100A9 undergoes nucleation-dependent polymerization as we have demonstrated previously and does not involve secondary nucleation.^12,33^ The kinetics of S100A9 fibrillation at different concentrations show that there is no noticeable lagphase (Fig. 3A), indicating that the protein misfolding and primary nucleation is a rate-limiting step. The global fit results in the values of critical nuclei size, *n_c_*,= 1.66 and combined rate constant *k_n_k_+_* = 2.05 104 mM^-1.66^ h^-2^.

**Figure 3.**
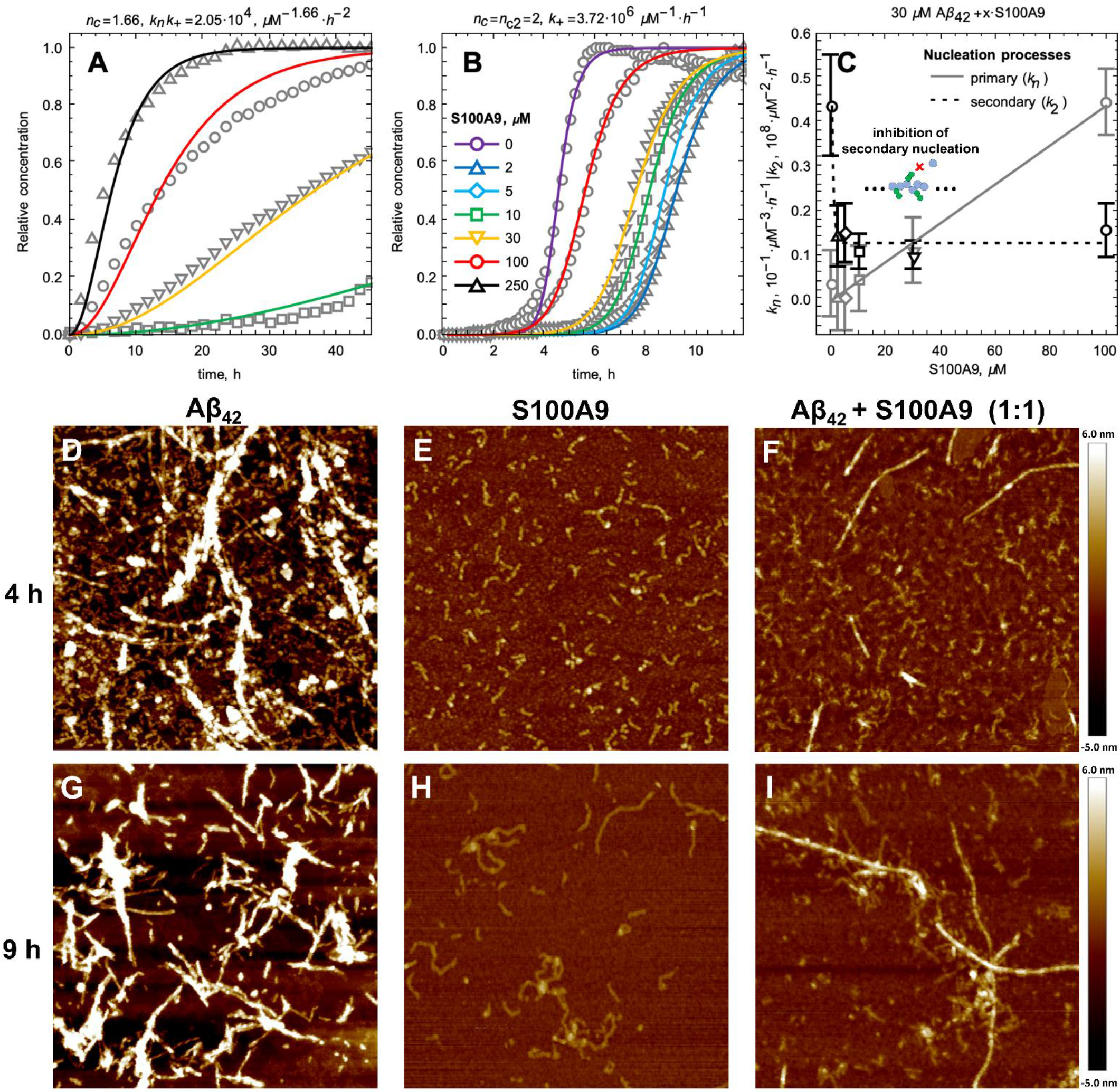
S100A9 affects Aβ_42_ primary and secondary nucleation pathways as revealed by the kinetic and AFM analyses. (A) Fibrillation kinetics of S100A9 monitored by ThT fluorescence and normalised to initial native S100A9 concentrations. Solid lines represent the global fitting by nucleation-dependent polymerisation model and experimental points are shown by grey symbols. (B) Normalised amyloid kinetics of Aβ_42_ and S100A9 mixture monitored by ThT fluorescence and fitted by using a secondary nucleation dominated model. Solid lines represent the fitting curves and experimental points are shown by grey symbols. 30 μM Aβ_42_, S100A9 concentrations are indicated in figure caption in corresponding colour coding (the same colour coding is used for A and B). Each solution contained 30 μM Aβ_42_ and added S100A9 concentration, respectively. (C) Dependences of the primary nucleation, *k_n_* (in solid grey), and secondary nucleation, *k_2_* (in dashed black), rate constants on S100A9 concentration as derived from the global fit presented in (B). AFM imaging of (D,G) Aβ_42_, (E,H) S100A9 and (F,I) Aβ_42_-S100A9 incubated for 4 h and 9 h, respectively, using 30 μM S100A9 and 30 μM Aβ_42_, in PBS, pH 7.4 and 42°C. Scan sizes are 2 x 2 μm.

The fibrillation curves of Aβ_42_-S100A9 co-aggregation display typical sigmoidal shape (Fig. 3B) characteristic for fibrillation of Aβ_42_.^11,32^ Incubation of 2 to 100 μM S100A9 alone manifested in significantly lower ThT signal, if any, compared to the signal of ThT bound to Aβ_42_ fibrils (Fig. S8A^†^ and B^†^). Therefore, in Aβ_42_-S100A9 mixture the major ThT signal arises from dye molecules bound to Aβ_42_ amyloids and those signals were used for fitting the fibrillation curves by the secondary nucleation dominated model as has been shown previously for Aβ_42_.^11,32^ The presence of S100A9 leads to increase of the lag-phase; the lowest 2 μM S100A9 concentration manifested in the most pronounced lag-phase increase to ca. 7 h, while 100 μM S100A9 results in ca. 4 h lag-phase (Fig. 3B). In the presence of increasing S100A9 concentration the ThT plateau level of Aβ_42_-S100A9 complexes decreases (Fig. S8A^†^). Most noticeable ThT signal decrease at highest S100A9 concentration in solution may reflect the coating effect of S100A9 species on the Aβ_42_ amyloid surfaces. In the fitting of Aβ_42_-S100A9 co-aggregation kinetics the elongation rate, *k_+_*, was set as a global fit parameter, i.e. shared for all fitted curves. The primary, *k_n_*, and secondary, *k_2_*, nucleation rates were set as fitting parameters, i.e. as variables for each of the fitted curves. Based on the reaction kinetic analysis we may conclude that the secondary nucleation rates for Aβ_42_-S100A9 complexes are significantly reduced (Fig. 3C). This is in agreement with the AFM observations of the increased length of Aβ_42_ carrier fibrils templating S100A9 amyloid on their surfaces compared to Aβ_42_ incubated alone (Fig. 1D, E and S2^†^). Interestingly, in the presence of 5 μM S100A9 in the mixture with Aβ_42_, the length of Aβ_42_ fibrils also increases, but to a smaller extend than in the presence of 30 μM S100A9; the median values of the corresponding length distributions are 0.68 μm *vs* 0.75 μm, respectively, as presented in Fig. S2^†^. The fibril length can be related to the rates of elongation and secondary nucleation^34^ and, assuming that the change in fibril length observed here is due to a change in secondary rate alone, we obtain the following approximation for the change in fibril length, *μ*,

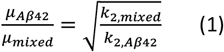

Given the change in length observed by AFM (Fig. 1D and E), we thus expect a decrease of the rate constant of secondary nucleation, *k_2_*, by approximately a factor of 3 in the presence of S100A9 compared to pure Aβ_42_. Indeed, a decrease of *k_2_* by approximately this value is also obtained from analysis of the aggregation kinetics (Fig. 3C), showing that these two orthogonal measurements yield consistent results. Thus, the blocking of Aβ_42_ fibrillar surfaces and its secondary nucleation by templating on them S100A9 fibrils leads to increase of their length.

By contrast, the distributions of lengths and heights of S100A9 fibrils remain the same (Fig. S2^†^) as whether they are fibrillated alone or in Aβ_42_-S100A9 mixture, including both S100A9 filaments templated on Aβ_42_ surfaces and free in solution. This indicates that as long as S100A9 fibrils were templated on Aβ_42_ fibril surfaces, their size distributions are not affected by Aβ_42_.

At the same time the higher rates of primary nucleation are consistent with the reduction of lag-phase of Aβ_42_-S100A9 co-aggregation in the presence of S100A9 (Fig. 3B and C). The hydrophobic properties of S100A9 dimers and their larger effective cross-sections compared to these of Aβ_42_ monomers may well serve also as nucleation sites for Aβ_42_, especially if S100A9 itself undergoes amyloid self-assembly.^12,33^ This implies that the effect of S100A9 on both primary and secondary nucleation of Aβ_42_ may depend on the degree of S100A9 polymerization.

AFM imaging was carried out in parallel to the amyloid kinetics to monitor amyloid development in time (Fig. 3D-I). After 4 h incubation Aβ_42_ alone self-assembles into a large amount of protofibrils and mature fibrils (Fig. 3D), S100A9 forms very short filaments (Fig. 3E) and Aβ_42_-S100A9 sample is characterized by both emerging Aβ_42_-like fibrils, though in significantly smaller quantity compared to Aβ_42_ incubated alone, and short protofilaments (Fig. 3F). After 9h, Aβ_42_ and S100A9 individually form their typical fibrils (Fig. 3G and H). By contrast, Aβ_42_-S100A9 sample displays the mature thick fibrils, characteristic for Aβ_42_, massively coated by distinct thin S100A9 filaments. They are present together with the short and thin filaments, characteristic for S100A9, in the surrounding solution (Fig. 3I), which is consistent with the corresponding images after 24 h incubation (Fig. 1C and S1^†^).

Further insights into the effect of non-aggregating S100A9 on Aβ_42_ amyloid fibrillation was provided by incubating both polypeptides at pH 3.0, where Aβ_42_ readily forms mature twisted fibrils (Fig. 4A and B), while S100A9 does not form amyloids at all (Fig. S9A^†^ and B^†^). In the presence of 3 μM non-aggregating S100A9, Aβ_42_ amyloid formation is delayed, as reflected in an increased lag-phase and decreased both growth phase slope and ThT plateau level (Fig. 4A). Aβ_42_ fibrillation is completely abolished in the presence of 30 μM S100A9, which is shown by both the absence of ThT signal and precipitation of unstructured aggregates observed by AFM imaging (Fig. 4C). The effects of the same concentrations of S100A9 fibrillar species on Aβ_42_ fibrillation is less pronounced, i.e. 3 μM S100A9 fibrillar seeds do not significantly perturb the Aβ_42_ fibrillation, while 30 μM seeds lead to some delay in amyloid formation and decrease in ThT fluorescence plateau (Fig. 4D). In the latter, Aβ_42_ fibrils are coated with S100A9 amyloid filaments as observed in AFM image (Fig. 4F). This indicates that the surfaces available for Aβ_42_ secondary nucleation are diminished by the presence of S100A9 amyloid coating, though S100A9 fibrillar species are less efficient in inhibiting Aβ_42_ fibrillation than non-aggregated S100A9.

**Figure 4.**
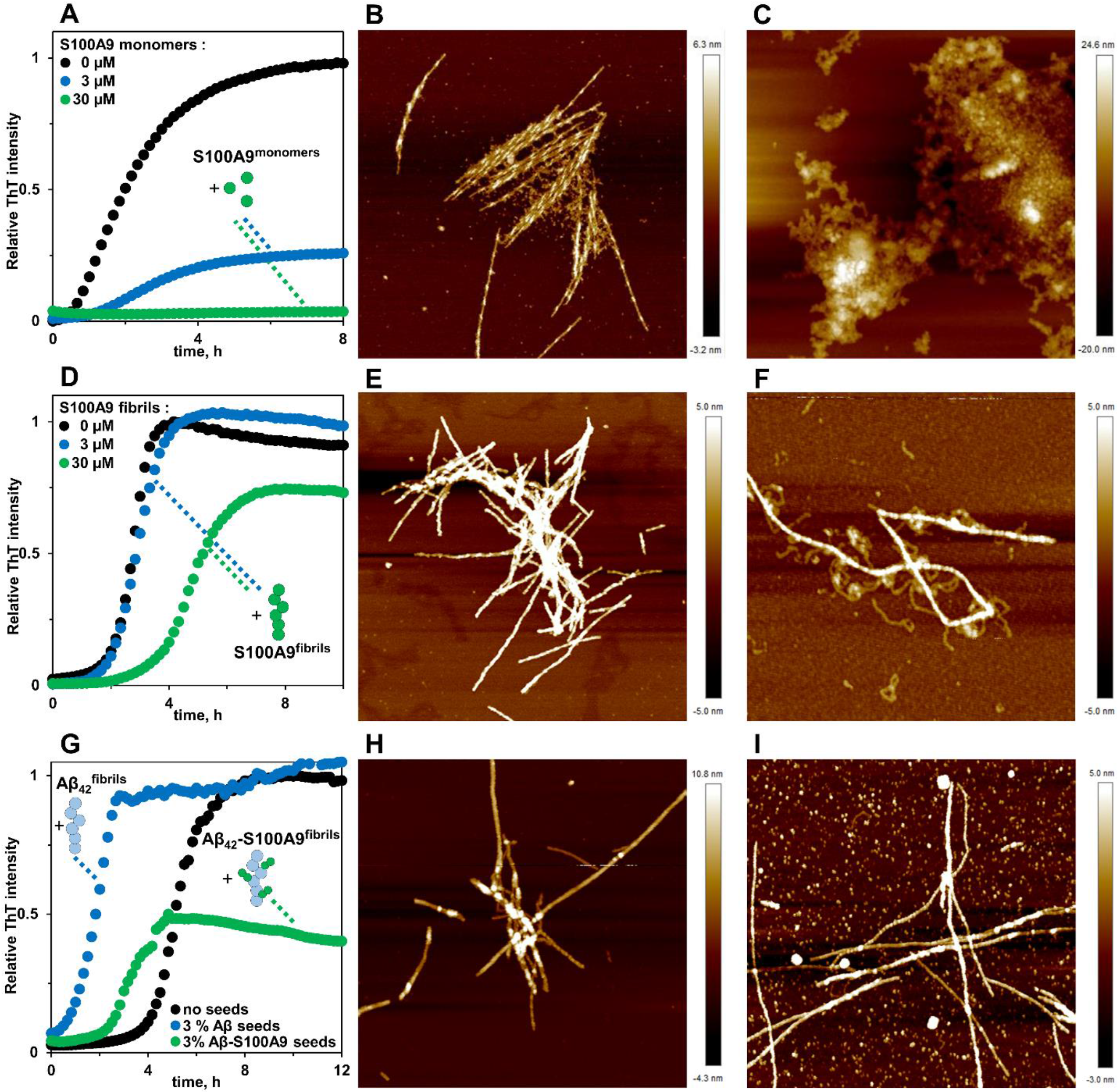
Non-aggregating and fibrillar S100A9 produce different effects on Aβ_42_ amyloid aggregation. (A) Fibrillation kinetics of the mixture of Aβ_42_ and S100A9 monitored by ThT fluorescence,10 mM HCl, pH 3, 42° C. Concentrations of added non-aggregating S100A9 are indicated in figure caption in corresponding colour coding. AFM images of (B) Aβ_42_ and (C) Aβ_42_-S100A9 (1:1 molar ratio) aggregates observed after 24 h incubation. (D) Fibrillation kinetics of Aβ_42_ in the absence and presence of S100A9 fibrillar seeds monitored by ThT fluorescence. Concentrations of added S100A9 fibrillar seeds are indicated in figure caption in corresponding colour coding. AFM images of (E) Aβ_42_ and (F) Aβ_42_-S100A9 (1:1 molar ratio) aggregates after 24 h incubation. (G) Fibrillation kinetics of Aβ_42_ in the absence and presence of the fibrillar seeds of Aβ_42_ and Aβ_42_-S100A9. Added fibrillar seeds are indicated in figure caption in corresponding colour coding. AFM images of (H) Aβ_42_ incubated with 3% Aβ_42_ and (I) Aβ_42_ with 3% Aβ_42_-S100A9 seeds after 30 h incubation. 30 μM Aβ_42_ was used in all experiments. Scan sizes are 2 x 2 μm.

Co-incubation of Aβ_42_ with 3% Aβ_42_ fibrils produces pronounced seeding effect on Aβ_42_ fibrillation, effectively abolishing the lag-phase and inducing mature fibril formation (Fig. 4G and H). By contrast, 3% Aβ_42_-S100A9 co-aggregates are much less efficient in shortening the lag-phase, while causing nearly twice decrease of ThT plateau level and leading to the formation of fibrils and round-shaped aggregates (Fig. 4G and I). Since under the seeding conditions the Aβ_42_ secondary nucleation pathways are dominant,^11^ the Aβ_42_-S100A9 seeds coated by S100A9 become less efficient than pure Aβ_42_ fibrils. Noteworthy, in the control experiments, we demonstrate that S100A9 amyloid kinetics are not affected either by cross-seeding with Aβ_42_ fibrils, even at 10% Aβ_42_ seeds, or seeding with S100A9 fibrils (Fig. S9^†^).

The binding of native S100A9 to Aβ_42_ fibrils was also examined by using a microfluidic diffusional sizing method as described in Materials and Methods^†^ and shown in Fig. 5A. The binding affinity of native S100A9 to Aβ_42_ fibrils was determined by measuring hydrodynamic radius, *R_h_*, in the presence of increasing Aβ_42_ fibril concentrations. The corresponding values of dissociation constant 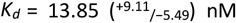 nM and stoichiometric ratio 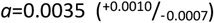 nM were determined from the resulting binding curve (Fig. 5B). Such stochiometric ratio corresponds to approximately one S100A9 binding site per ca. 300 Aβ_42_ monomers in the Aβ_42_ fibril. Based on the calculation of the number of monomers per unit of the Aβ_42_ fibril length derived from the cryo-electron microscopy,^35^ the distance between S100A9 binding sites on Aβ_42_ fibril would be ca. 100 nm. AFM analysis indicates that the binding sites of S100A9 on Aβ_42_ fibrils can be visualized with an average distance of ca. 100 nm between S100A9 filaments templated on Aβ_42_ fibril in Aβ_42_-S100A9 complexes (Fig. 5C and S10^†^). These numbers are broadly consistent with the stoichiometry determined by microfluidic diffusional sizing. Thus, by two independent methods we have demonstrated that the distance between the S100A9 binding and secondary nucleation sites on Aβ_42_-S100A9 is about the same.

**Figure 5.**
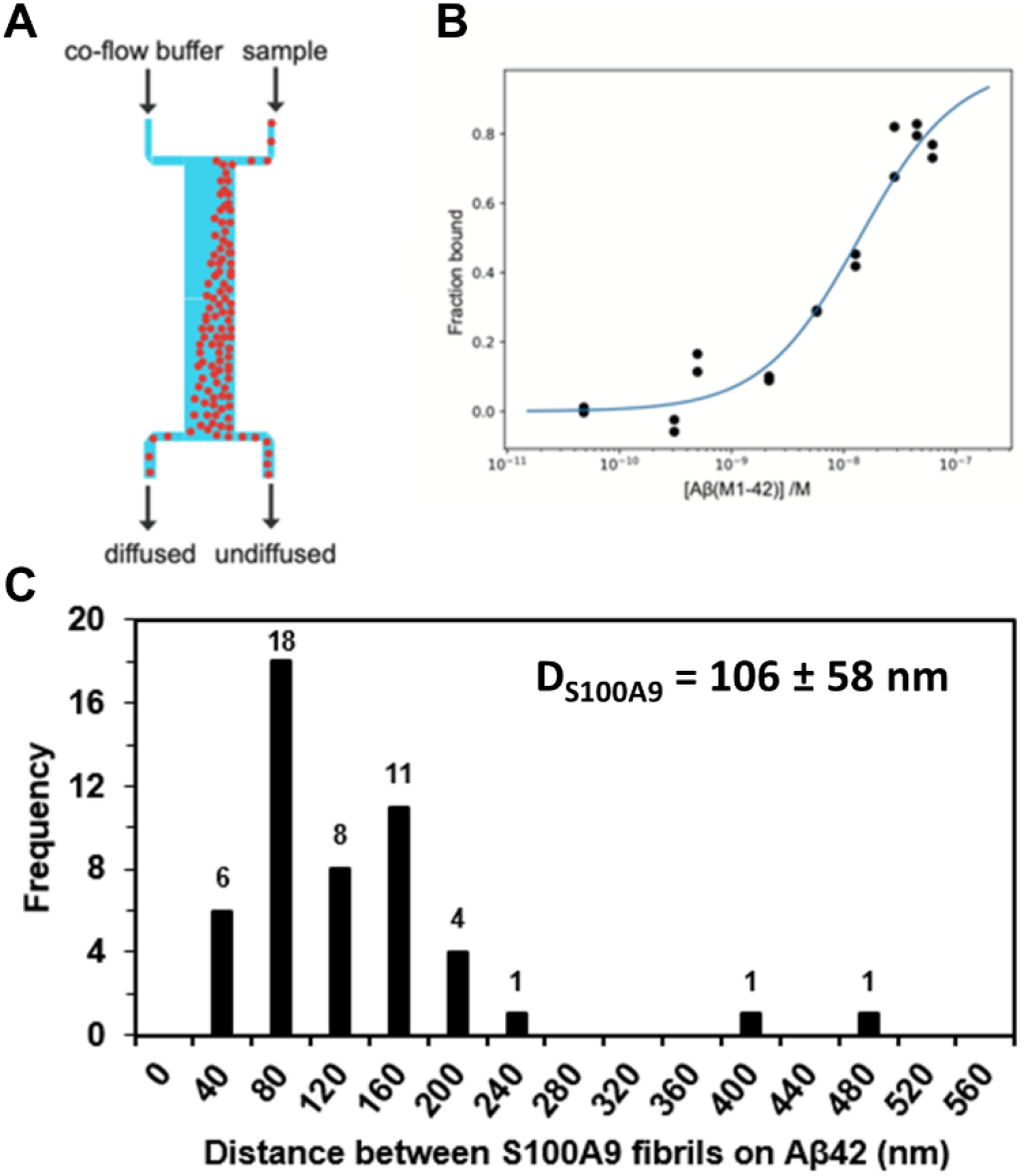
Native S100A9 binding to A β42 fibrils measured by microfluidic diffusional sizing. (A) Scheme of a microfluidic channel used to measure binding affinity between native S100A9 and Aβ_42_ fibrils (Materials and Methods^†^). (B) Binding curve for the interaction between S100A9 and Aβ_42_ fibrils, from which dissociation constant and stoichiometric ratio were determined by Bayesian analysis. (C) Distribution of the distances between S100A9 fibrils templated on the surface of Aβ_42_-S100A9 amyloids imaged by AFM. 30 μM of each polypeptide were co-incubated for 24 h in PBS, pH 7.4 and 42°C.

## Conclusions

Inflammation is central to the onset of AD and many other neurodegenerative diseases, however the mechanistic insights into how inflammatory events are linked to the amyloid formation remain unclear. By using CDMS in combination with microscopy, kinetic analysis and microfluidic binding assay we have demonstrated that proinflammatory S100A9 protein, involved in AD and range of other neuroinflammatory and neurodegenerative diseases, co-assembles with Aβ fibrils, forming a new type of hetero-amyloid complexes.^9,10,33^ In these complexes the autocatalytic surfaces of Aβ_42_ fibrils template S100A9 amyloids, where each component represents a homo-molecular domain in the hetero-molecular Aβ_42_-S100A9 co-assembly. These change the dynamics of Aβ amyloid aggregation and distribution of sizes of resulting co-assembled Aβ_42_-S100A9 complexes. The formation of larger Aβ_42_-S100A9 complexes may sequestrate smaller and more toxic species from the environment, which is consistent with our previous finding that co-aggregation of S100A9 with either Aβ_42_ or Aβ^40^ mitigate the overall amyloid cytotoxicity.^9^ Thus, these findings contribute to understanding of amyloid co-aggregation processes both from a fundamental perspective and in revealing disease relevant processes.

Here we also exemplified analytical methods applied in synergy for the accurate analysis of such complex system as amyloid co-aggregation. We provide an analytical framework to utilize the capacity of CDMS, which can directly distinguish highly heterogeneous populations of amyloid co-aggregates in two dimensions, in combination with other biophysical techniques, assessing the bulk ensemble of molecular species, which together enable us to discriminate the amyloids originated from homo and hetero-molecular co-aggregation reactions. We have demonstrated the broad consistency in quantitative and qualitative measurements produced by those complementary techniques – CDMS and AFM (hetero-assemblies were observed by both methods); AFM and kinetic analysis (revealing the correlation between the fibrillar length and reduced secondary nucleation rates in the hetero-complexes) as well as microfluidic binding assay and AFM (consistency in stoichiometry of binding/templating of S100A9 on Aβ_42_ fibrils).

The genuine understanding of the mechanisms underlying Aβ_42_ and S100A9 driven amyloid-neuroinflammatory cascade in AD may also provide prospective target for therapeutic interventions and lead to the development of therapy for a cureless disease as the current approaches to target only one protein type did not mature.

## Conflicts of interest

There are no conflicts to declare.

## Acknowledgements

We thank Daphna Fenel and Dr Guy Schoehn, from the IBS/UVHCI platform of the Partnership for Structural Biology in Grenoble (PSB/IBS) for the Electron Microscopy. We acknowledge financial support from VR-M and the Medical Faculty, Umeå University (LAM-R), for RA and HK the project STIM – REI, Contract Number: KK.01.1.1.01.0003, funded by the European Union through the European Regional Development Fund – the Operational Programme Competitiveness and Cohesion 2014-2020 (KK.01.1.1.01); from the BBRSC (TPJK), the ERC PhyProt (agreement no. 337969) (MMS,TS,CKX, GM, TPJK), the Frances and Augustus Newman Foundation (TPJK) and the Centre for Misfolding diseases (MMS, TS, CKX, GM, TPJK).

## Materials and methods

### Amyloid fibril formation

0.5 mg of freeze-dried Aβ_42_ peptide (American Peptide Company) was freshly dissolved in distilled water at pH 11.0 and filtered through a 0.22 μm spin membrane filter (Millex, ref. SLGV013SL) to remove any aggregated species. Aβ_42_ fibrils were prepared by incubating 100 μM Aβ_42_ peptide in phosphate buffer saline (PBS, Medicago) at pH 7.4 and 42°C. Aβ_42_ was also subjected to amyloid aggregation in 10 mM HCl buffer at pH 3 and 42°C.

S100A9 protein was expressed in *E.coli* and purified as described previously.^1^ Freeze-dried S100A9 was dissolved on ice in PBS buffer at pH 7.4 and 42°C to 400 μM concentration. Before incubation, S100A9 samples were filtered through a 0.22 μm spin membrane filter to remove any aggregates. To conduct experiments at pH 3.0 freeze-dried S100A9 protein was dissolved directly into 10 mM HCl buffer at pH 3 and 42°C.

### ThT fluorescence assay

ThT dye is known to bind specifically to β-sheet containing amyloids, and thus enables to follow the kinetics of amyloid self-assembly.^2^ Aβ_42_, S100A9 and mixed solutions of Aβ_42_ and S100A9 were prepared at different concentrations on ice, transferred into 96-well plates and then 20 μM ThT was added to each well. The plates were immediately covered, placed into a Tecan F200 PRO plate reader and incubated at 42°C by using 432 rpm orbital shaking every 10 min. ThT fluorescence was also recorded each 10 min, using 450 nm and 490 nm for excitation and emission, respectively. Each sample was incubated in triplicates.

### Reaction kinetics fitting

The aggregation kinetics of Aβ_42_-S100A9 co-aggregates and S100A9 were fitted by Amylofit software.^3^ Secondary nucleation dominated model was used for Aβ_42_-S100A9 with elongation rate set as a global fitting parameter and critical nuclei size for primary and secondary nucleation was set to 2 (*n_c_* = *n_2_* = 2). Number of basin hops was set to five. By using described parameters, the fitting was performed seven times in order to estimate the accuracy of the fitted values, which is illustrated by the error bars in the corresponding figure. In the fitting of the co-aggregation kinetics, the ThT signal from S100A9 fibrils was neglected, since the ratio of the plateau intensities for the similar concentrations of S100A9 and Aβ_42_ is much less than 1 (I_S100A9_/I_Aβ_ ≪ 1). Nuclear-dependent polymerisation model was used to fit the S100A9 kinetics. Critical nucleus size, *nc*, and combined rate constant, *k_n_k_+_*, were used as global fit parameters.

### AFM imaging

15 μL of each sample were deposited on the surface of mica for 30 min, washed 5 times with 200 μL deionized water and left to dry overnight at room temperature. AFM imaging was performed in a PeakForce QNM mode in air by using a BioScope Catalyst atomic force microscope (Bruker). Resolution was set at 512 x 512 pixels, scan rate was 0.51 Hz, and scan sizes were 2 x 2 and 5 x 5 μm. Bruker RTESPA and SNL cantilevers were used. Heights of amyloid fibrils were measured in the cross-sections by using a Bruker Nanoscope analysis software, while their lengths were measured by using ImageJ software.

### AFM size distributions

Histograms of AFM fibril length and cross-sectional height distributions were built up by using the kernel density estimate and resampling techniques. Each distribution was built up by following procedures: firstly, the empirical distribution function was constructed for each observed AFM data set by using Gaussian kernel with bandwidth selected by Wand’s method;^4^ secondly, the sample size of 10^4^ was drawn from each distribution function. Thirdly, the histogram with bin size equal to the above bandwidth was constructed for each data set. This procedure was applied to all experimental data sets. The above resampling procedures did not affect the distribution parameters such as median and its deviation.

### CDMS instrument

The CDMS measurements were performed on home-built CDMS instrument in the single pass mode and the RMS detector noise was ~100e. In CDMS, an ion passes through a metal tube. A positive ion entering an isolated conducting tube induces a negative charge on the inner surface and a positive charge on the outside. The induced charges are maintained until the ion exits, at which point they dissipate. The mass-to-charge ratio (*m/z*) of the trapped ion is determined from the time-of-flight (TOF) *Δt*, *i.e.* time delay between the positive and negative pulses that corresponds to the entrance and the exit from the detector tube.

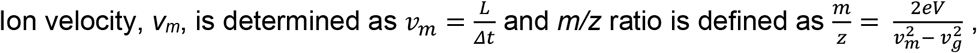

where *L = 3.75 cm* is length of the detector tube, *eV=155 V* – electrostatic acceleration voltage and *V_g_= 375 m/s* – the ion velocity due to the free gas expansion. Indeed, in our system a correction is needed to take into account the initial kinetic energy imposed on the ion by the free jet expansion of the gas prior to acceleration by the electric field.

### Electrospray ionization (ESI) conditions

To enhance ionization efficiencies, 25% methanol was added to all samples of corresponding amyloid fibrils and 22.5 μM final polypeptide concentrations were reached prior injection into the ESI source. The amyloid samples were injected at flow rates of typically 0.2-0.6 mL/h and entered the electrospray chamber through a 0.1 mm internal diameter stainless steel capillary tube located inside the needle tip. Nitrogen gas was injected between the end cap and the transfer glass capillary and was flown through a heater, typically set at 200°C. The vacuum interface was composed of a glass transfer capillary that passes the ions into the first stage of the vacuum system, an end cap, a skimmer between the first and second vacuum stages, a hexapole ion guide and an exit lens. The ESI source generates charged macroions, which are guided by an ionic train to the mass spectrometer. Ions are guided up to a vacuum stage chamber (~5.10^−6^ mbar) and directed through the charge detection device. Importantly, the low charged population of amyloid fibrils was observed here due to improved signal to noise ratio of the home-built CDMS instrument. In our previous studies on amyloid populations we were able to detect only ions with charges higher than ~300 e.^11,12^ In the present work, the limit of charges was significantly reduced to ~200 e, due to improvements in the noise level in the pick-up signal and the addition of frequency filters.

### CDMS data processing

We used a home-developed Windows-based software to record chromatograms – VISUAL C++. The program calculates the time between the maxima of the positive and negative pulses, the amplitudes of two pulses and the ratio between their absolute values. A high-frequency filter is added to the data processing of traces in order to remove peak artefacts. Residual droplets are excluded by using post-processing thresholds for TOF (> 95 μs). In this work, only ions with charges higher than ~200 e that both enter and exit the tube, are counted. Events for which the absolute values of the amplitude ratios between the first and the second pulses are greater than 1.5 or less than 0.75 are automatically excluded. Finally, the corresponding ion counting rate ranges around 50 ions/s. For each ion the mass is deduced from its *m/z* and *z* values. For each sample, measured ions were filtered with respect to molecular mass (< 1 GDa) and charge (< 3 ke). Resulting sample sizes used in this study were thus 2863 for Aβ_42_, 839 for S100A9 and 1771 for Aβ_42_-S100A9 sample.

### Ranking of amyloid particles according to their CDMS charge to mass ratio (z/m)

The values of CDMS *z/m* ratios for each individual ion within each amyloid data set were arranged in descending order. Following the ranking, the positions of individual charged ions were plotted along *x*-axis, while *z/m* ratios were shown along *y*-axis. These values were compared to the *z/m* ratios of Aβ_42_ and S100A9 monomers, calculated based on their amino acid sequences at pH 7.4 by using a ProteinCalculator v.3.4 (protcalc.sourceforge.net).

### CDMS data analysis

In order to analyse the data in two dimensions of mass and charge, the joint mass-charge frequency distributions were constructed as shown schematically in Fig. S1 and described below. Firstly, the range of data was binned with respect to charge and mass; secondly, the numbers of ions falling into corresponding bins were counted and thirdly, the counts were divided by the sample size, i.e. total number of ions in the sample. The optimal bin width was calculated for each molecular mass distribution using one level recursive Wand method.^4^ Optimal bin width ensures that the histogram is not oversmoothed. The same bin number was used for all samples, i.e. 50 bins. The same number of 50 bins was used for the charge range since molecular mass and charge are dependent quantities.^5^

We have constructed three joint frequency distributions with respect to mass and charge termed as follows: Φ(*m,z*) = Φ_*Aβ*_42__, Φ_S100A9_ and Φ_*Aβ*_42_−S100A9_. Subsequently, difference frequency distributions were constructed by pointwise subtraction:

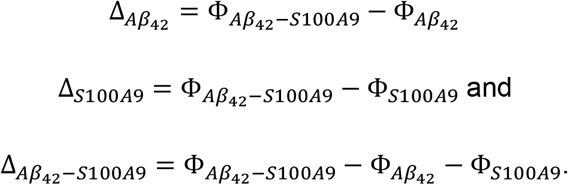

Since the range of *Φ* and *Δ* spans more than two orders of magnitude, the following logarithmic transformation 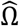 was used:

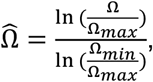

where: *Ω* denotes either of the following *Φ* or *Δ* distributions, *Ω_min_,Ω_max_* – minimal and maximal values among three distributions of either Φ_*Aβ*_42__, Φ_S100A9_ and Φ_Aβ_42__–S_100A9_ or Δ_Aβ_42__, Δ_S100A9_ and Δ_Aβ_42__–_S100A9_, respectively. Taking the minimal and maximal values across all three *Φ* or *Δ* distributions, respectively, ensures that after the log-transformation samples can be compared. The ranges of the difference distributions *Δ* are not symmetrical around 0, therefore two log-transformation have been performed separately for negative and positive values. In order to keep scaling symmetrical around 0, the widest range of all Δ was used to select minimal and maximal values for log-transformation. The absolute values were used for log-transformation and minus was assigned for the negative range of the log-transformed *Δ* values (Fig. S1).

The log-transformed distributions of *Φ* and *Δ* are represented by maps, where the colour corresponds to the frequency value. In order to keep the perception of the colour-mapped scalar value (frequency) linear with respect to human eye colour perception, the diverging colour scheme suggested by Moreland^6^ was used. It was scaled between 0 to 1 for the log-transformed frequency distributions *Φ* and between −1 to 1 for the log-transformed difference distributions *Δ.* We term the corresponding log-transformed frequency distributions as *frequency maps* and log-transformed difference frequency distributions as *difference maps*, respectively (Fig. S1).

### Simulated CDMS distribution

Probability mass function (PMF) was used to characterize the distribution of a discrete random variable such as mass. It associates the probability to any given number that the random variable will be equal to that number. We have simulated the molecular mass distribution of the fibril mixture based on CDMS molecular mass distributions of Aβ_42_ and S100A9 individual samples. In the case of separately formed fibrils the samples can be considered independent in mathematical sense, i.e. PMF of Aβ_42_ fibrillar sample does not depend on the PMF of S100A9 amyloid sample, which is not the case when co-aggregation occurs, i.e. for Aβ_42_-S100A9 amyloid sample. Imposing additional condition that sticking probabilities of fibrils do not depend on the fibril size and their type, we have calculated the distribution of molecular mass of preformed fibril mixture. Then PMF of mixed fibrils is the convolution of PMFs of individual components as follows: if *X* = Aβ_42_, *Y* = S100X9 and *Z* = *X* + *Y* = Aβ_42_+S100A9, then:

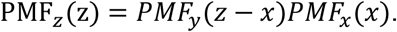

Mathematica 12 software was used to make statistical calculations and visualisation of results.

### Microfluidic binding assay

The microfluidic binding assay was used here as reported previously.^8–10^ Stream of fluorescently labelled analyte was injected from one side in a microfluidic channel alongside with auxiliary buffer flow coming from the opposite side into the channel under laminar flow conditions, allowing mixing by diffusion only (Fig. 5A). By tracking the spatial diffusion of the analyte into the co-flow buffer, the hydrodynamic radius, *R_h_*, was determined. The binding affinity, *K_d_*, and stoichiometry, *α*, were derived from the changes in *R_h_* by using Bayesian analysis.

The S100A9 protein was fluorescently labelled by incubation with three molar equivalents of Alexa Fluor 647 (ThermoFisher, UK) for 1 h after buffer exchange to NaHCO3 (0.1 M, pH 8.3). The protein conjugate was purified on a Superdex 200 increase 10/30 gL column (GE Healthcare, US) with a flow rate of 0.5 mL/min with PBS as elution buffer, containing 0.02 % NaN_3_ (w/v).

Aβ_42_ monomer was purified as reported previously, including a series of purification cycles.^7^ For the final purification step monomeric Aβ_42_ was incubated in Gdn-HCl (8 M in sodium phosphate buffer) for 1 h and purified on a Superdex 75 increase 10/30 gL column with sodium phosphate buffer (20 mM, pH 8.0, with 0.2 mM EDTA). Aβ_42_ fibrils were obtained by incubating 30 μM monomeric Aβ_42_ in 20 mM sodium phosphate buffer (pH 8.0, containing 0.2 mM EDTA) at 37°C with double orbital rotation (400 rpm) in a 96-well plate in a FLUOstar OPTIMA plate reader (BMG Labtech). The aggregation of Aβ_42_ was followed by measuring the fluorescence increase of 20 mM ThT in one similar aliquot. After completion of the aggregation reaction, the fibrils were collected and used in the microfluidic binding assay.

S100A9 was incubated with varying concentration of Aβ_42_ in sodium phosphate buffer and Tween 20 (0.02 % v/v) for 2 h. The increase in size upon binding was detected by applying microfluidic diffusional sizing in Fluidity One W (Fluidic Analytics, Cambridge, UK) series of purification cycles. For the S100A9 protein in absence of Aβ_42_ fibrils a hydrodynamic radius *Rh* = 2.52 ± 0.13 nm was determined, which is in a good agreement with the theoretically expected radius of 2.46 nm for a dimer of 13,242 Da protein.

### Analysis of microfluidic binding data

In our binding equilibrium we have S100A9 (A) and fibrillar Aβ peptide (B):

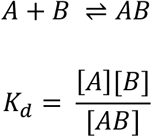

Denoting the total concentrations of A and B as [*A*]_0_ and [*B*]_0_, respectively, we obtain the following expression for the concentration of A bound to B, [*AB*]:

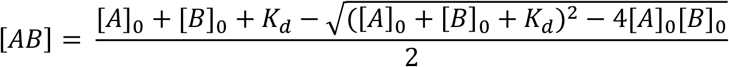

It is important to note that B here denotes the concentration of fibril binding sites, rather than monomers in the fibril. The hydrodynamic radius is calculated from measurements of the fluorescence intensities of the ‘diffused’ and ‘undiffused’ channels, termed *I_d_* and *I_u_*, respectively. However, unlike the radius, the fraction of labelled substrate that ends up in the ‘diffused’ channel, 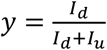, is easy to relate to the concentration of A bound. We thus describe the analysis here in terms of *y*, while the conversion to *R_h_* can be performed after fitting. We also define the parameters and *r_b_*, denoting the fractions of free and bound labelled substrate, respectively, that are detected in the ‘diffused’ channel. The following equations describe the intensities in each channel:

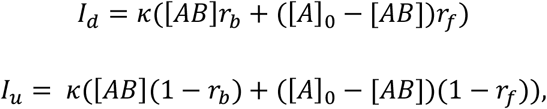

where *κ* is a constant relating label concentration with fluorescence intensity detected.

We can therefore express the predicted fraction in the diffused channel by:

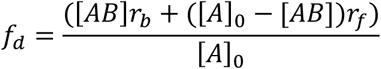

with [AB] determined by the equation above. Therefore, the predicted fraction in the diffused channel is a function of *r_f_*, *r_b_*, *K_d_*, [*A*]_0_, and [*B*]_0_, thus denoted by *f_d_*(*r_f_*, *r_b_*, *K_d_* [*A*]_0_, [*B*]_0_.

Finally, the stoichiometry, *α*, of the Aβ fibril : S100A9 (B : A) interactions are also not known. In order to account for this, we define *α* as the number of S100A9 binding sites per Aβ monomer within the Aβ fibrils, so that [*B*]_0_ = *α*[*B*]_*tot*_ where [*B*]_0_ represents the concentration of binding sites, and [*B*]_*tot*_ the fibril concentration in terms of monomer equivalents. We assume that the radii of singly or multiply bound fibrils are equal, and that there is no cooperativity in the binding. We therefore have the following expression for [*AB*]:

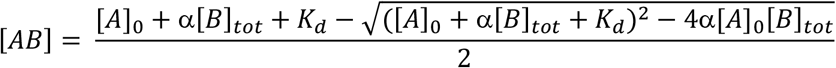

The expression for the predicted fraction to end up in the diffused channel is therefore *f_d_*(*r_f_*, *r_b_*, *K_d_*, *α* [*A*]_0_, [*B*]_0_), where there are 4 unknown parameters, *α*, *r_f_*, *r_b_*, and *K_d_.* Analysis then proceeds via Bayesian inference. For and *r_b_* we assume the prior is flat in linear space, whereas for the *K_d_* and a a prior, that is flat in logarithmic space, is chosen.

We assume our experimental measurement data to be normally distributed about the true value; our likelihood function is therefore a Gaussian, centered on the theoretical measurement value.

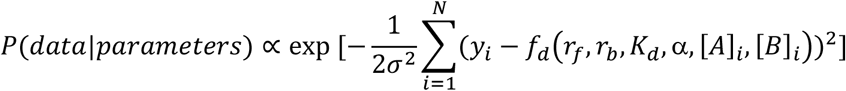

where *f_d_* is defined as above, [*A*]_*i*_ and [*B*]_*i*_ are the concentrations of S100A9 binding sites and Aβ, respectively, in the *i^th^* measurement, and *y_i_* is the fraction of diffused labelled component in the *i^th^* measurement. In order to define an appropriate standard deviation, *σ*, for each dataset, we calculate the standard deviations of repeats of each measurement and use the maximum of these values as a global standard deviation for that dataset. Inference is performed by calculation of the 4-dimensional posterior at evenly spaced points for all parameters, at 50 points each for *η* and *r_b_*, and 100 points each for *α* and *K_d_*, with ranges chosen such that there is no clear probability mass outside the sampled region. Marginalisation is used to obtain 1-dimensional distributions and thus calculate errors for the parameters.

## Author contribution

J.P., I.A.I. and L.A.M.-R. conceived and designed the work, and wrote the manuscript. J.P. and L.O. performed the *in vitro* characterizations of the amyloid fibrillation. H.F. and R.A. conceived, performed and analysed CDMS experiments. I.A.I. conducted conceptualization, formal analysis, methodology, validation and visualisation. M.M.S. and T.S. performed microfluidic experiments; the binding and kinetic analyses were conducted by C.K.X., G.M. and T.P.J.K. M.M., G.M. and V.S. prepared polypeptide samples. J.P., I.A.I., E.G. and L.A.M.-R. coordinated all experiments and compiled the results. All co-authors discussed and commented on the manuscript

## Supplementary figures

**Figure S1.**
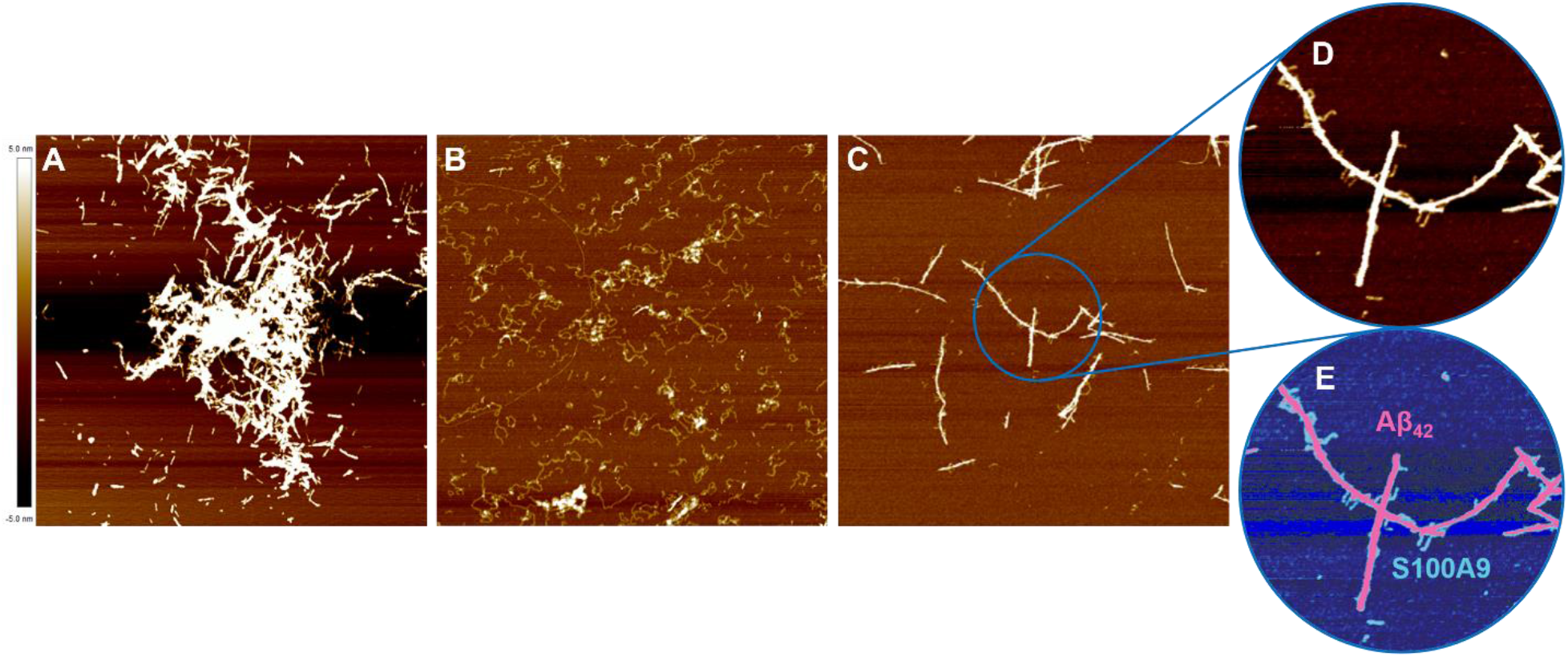
AFM analysis of Aβ_42_, S100A9 and co-aggregated Aβ_42_-S100A9 amyloids. AFM images of amyloids of (A) 30 μM Aβ_42_, (B) 30 μM S100A9 and (C) Aβ_42_-S100A9 (30 μM Aβ_42_ and 5 μM S100A9 were co-incubated). Amyloids were formed after 24 h incubation in PBS, pH 7.4, 42°C. Scan sizes are 5 x 5 μm. (D,E) Magnified images of Aβ_42_ fibrils with S100A9 amyloids templated on their surfaces presented in arbitrary colour scheme.

**Figure S2.**
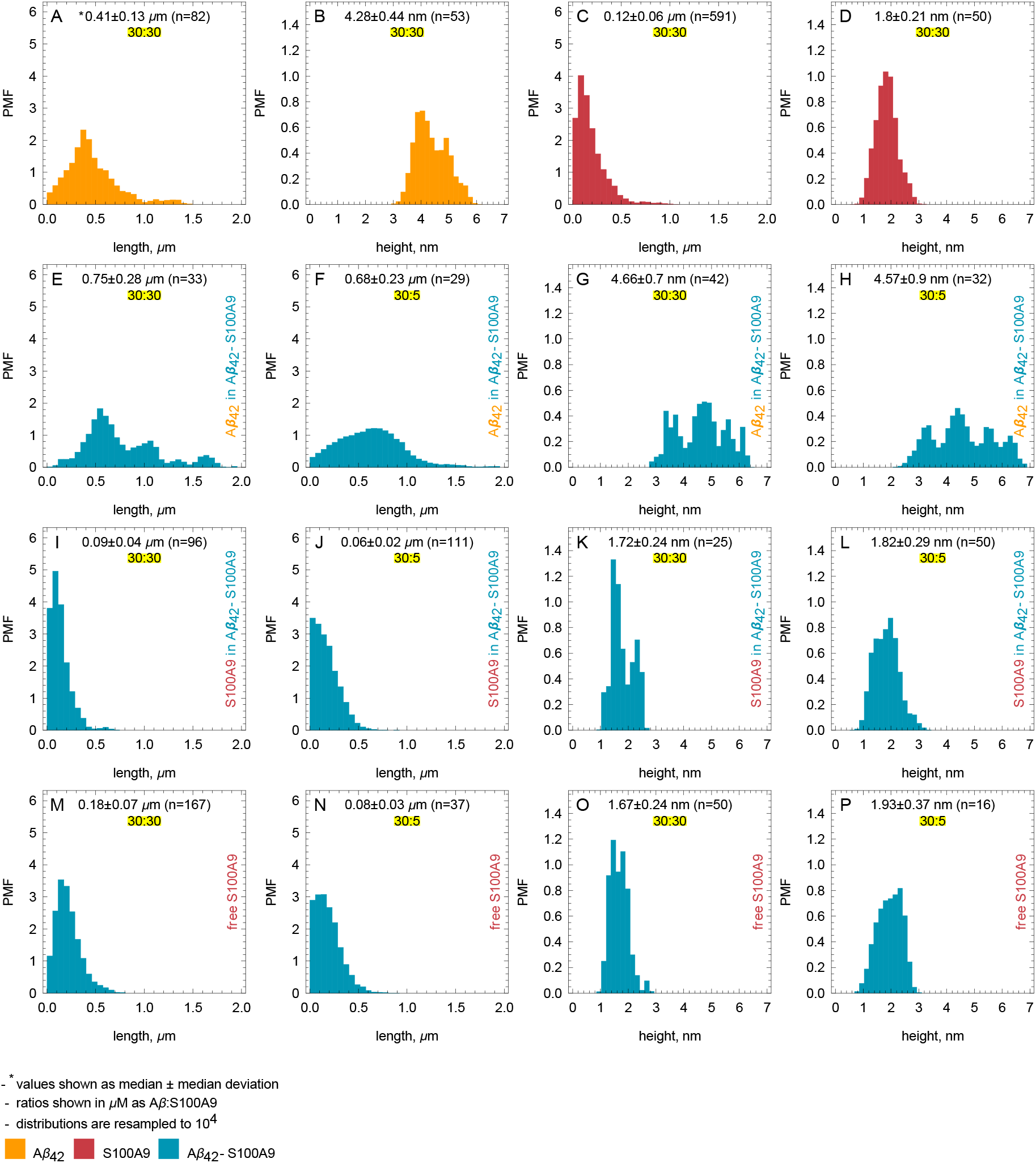
Length and height distributions of amyloid fibrils measured by AFM. Length of fibrils were measured by ImageJ software. The heights of amyloid fibrils were measured by Bruker Nanoscope analysis software in the AFM cross-sections. Probability mass function (PMF) defined as probability of finding fibril with specific length or height is indicated along *y*-axes. The measured values of lengths or heights are indicated along *x-*axes. The length and height distributions for Aβ_42_ fibrils are shown in yellow, for S100A9 fibrils – in red and for Aβ_42_-S100A9 amyloids – in blue. The distributions for Aβ_42_ fibrils within Aβ_42_-S100A9 complexes are shown in 3^rd^ row; for thin S100A9 fibrils templated on the surfaces of Aβ_42_-S100A9 complexes – in 4^th^ row and for free S100A9 in the Aβ_42_-S100A9 samples in 5^th^ row. The molar ratios of Aβ_42_ and S100A9 co-incubated in Aβ_42_-S100A9 samples are shown in each figure legends as 30:30 or 30:5, respectively. *The median values of corresponding distributions and sample sizes are also shown in each figure. The distributions are resampled to 10^4^ (See ESI Material and Methods).

**Figure S3.**
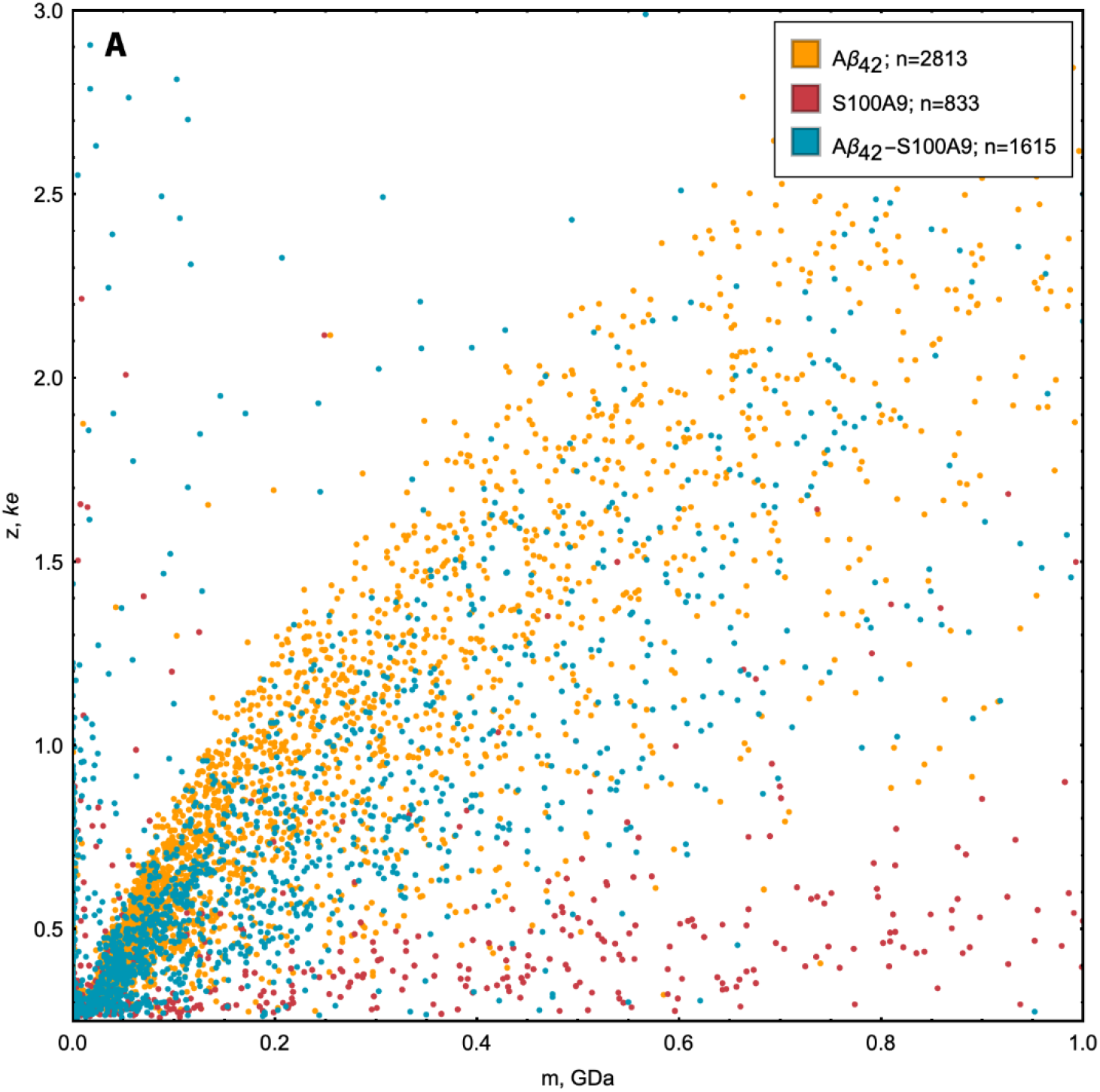
CDMS analysis of the amyloid samples. CDMS populations of amyloid particles with corresponding molecular masses (*m*) and charges (*z*) for the amyloid samples of Aβ_42_ (in yellow), S100A9 (in red) and Aβ_42_-S100A9 (in blue), respectively.

**Fig. S4.**
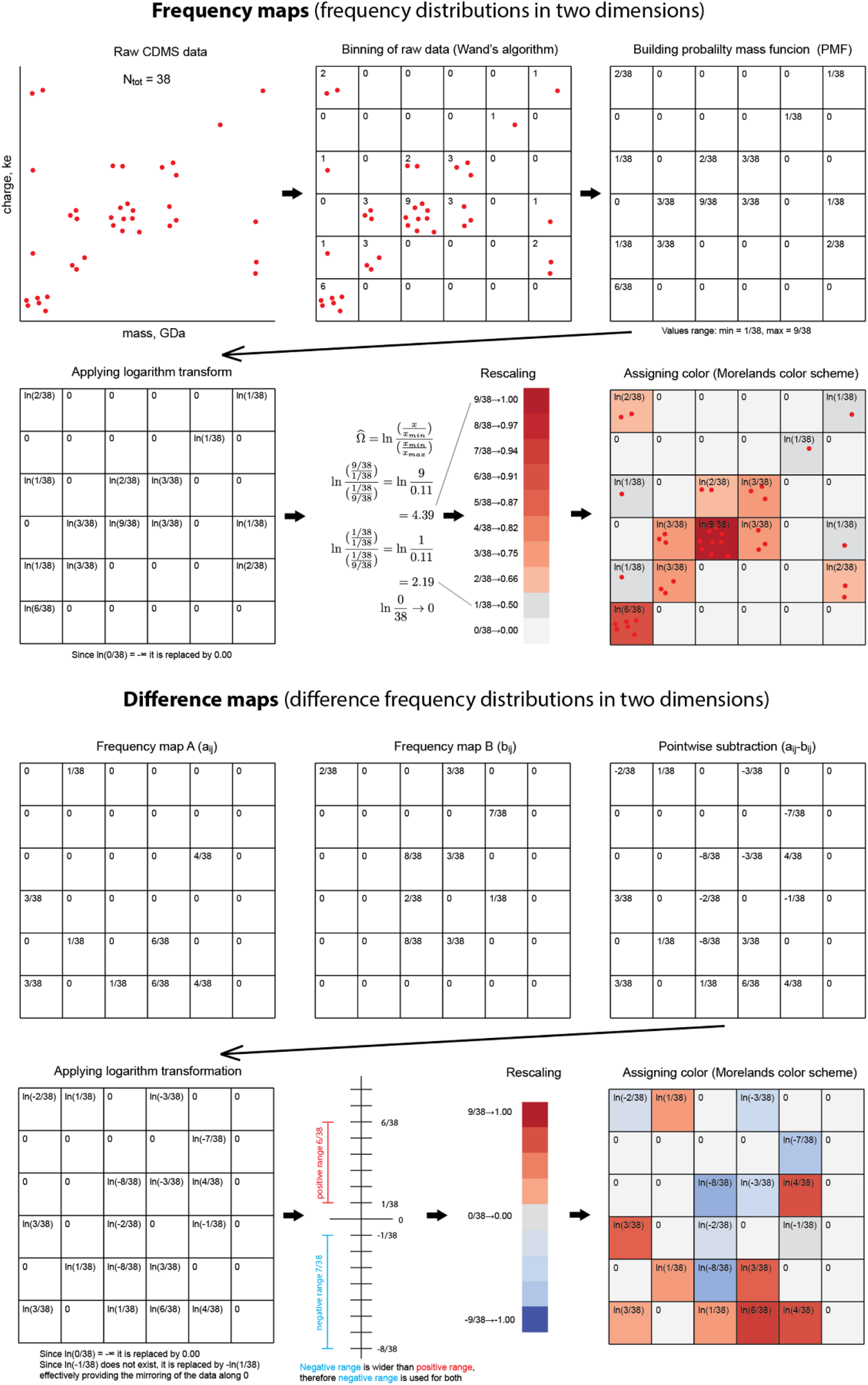
Schematic presentation of the sequence of steps involved in calculation the CDMS frequency and differential maps, respectively.

**Figure S5.**
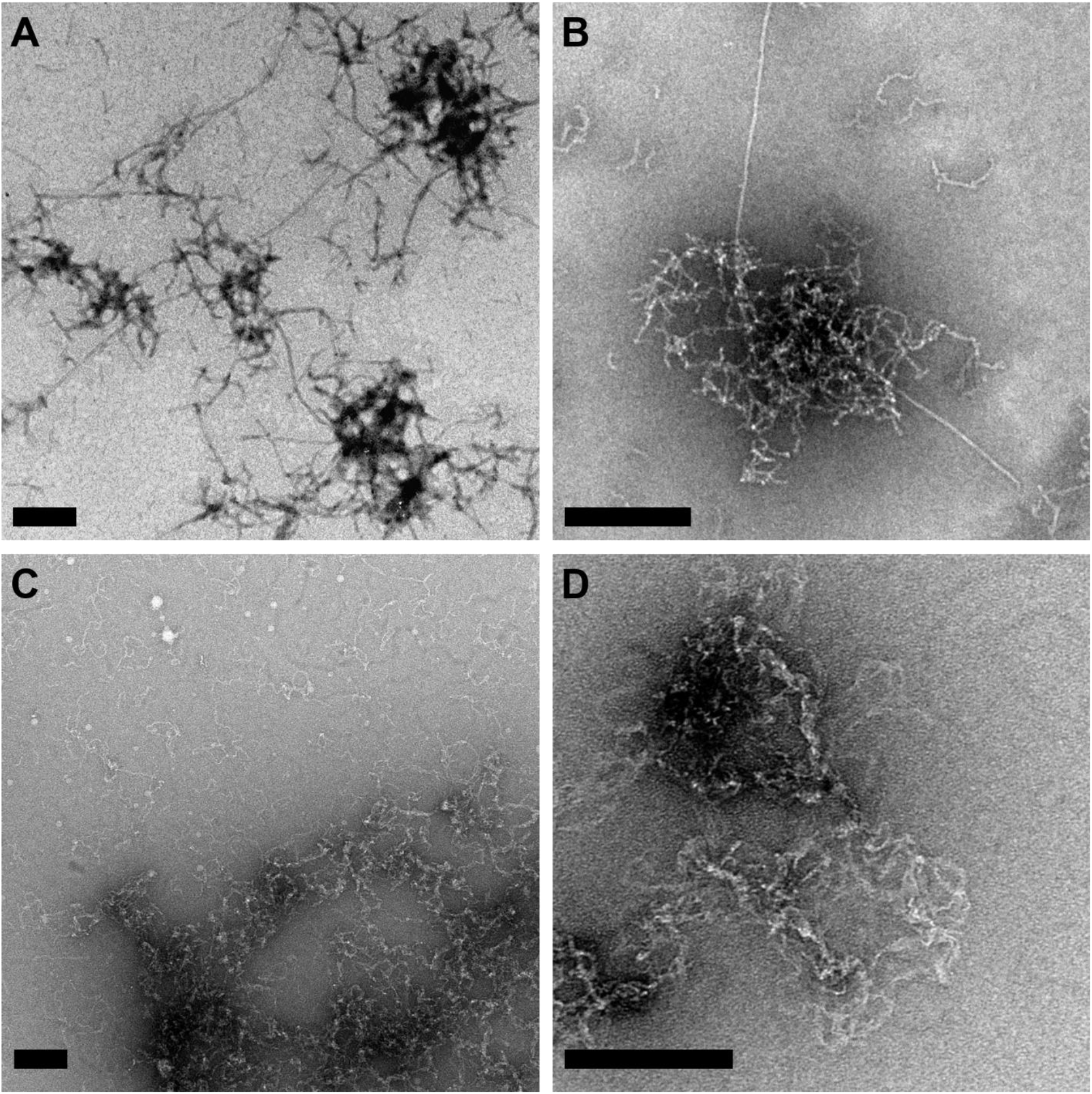
TEM images of Aβ_42_ and S100A9 amyloids. (A,B) TEM images of Aβ_42_ fibrils and their clusters presented at two magnifications. (C,D) TEM images of S100A9 amyloid fibrils and their coiling into amyloid clumps presented at two magnifications. Scale bars are 200 nm.

**Figure S6.**
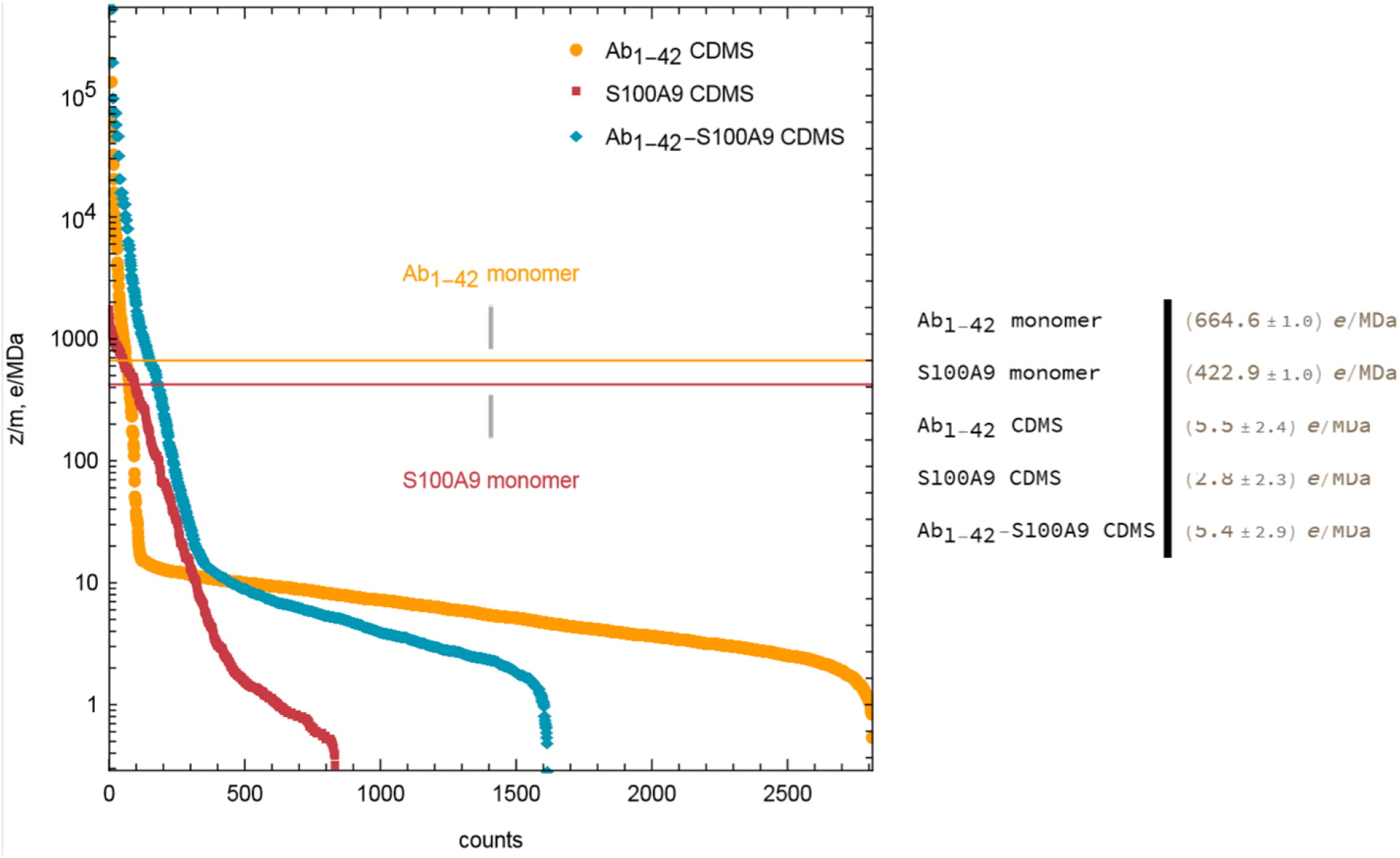
Ranking of amyloid particles according to their charge to mass ratio (*z/m*). Ratios of *z/m* of amyloid particles were taken from CDMS data and shown along *y*-axis. Positions of individual charged ions after their ranking are shown along *x*-axis. The ion positions corresponding to Aβ_42_ fibrils are shown in yellow, for S100A9 fibrils – in red and for Aβ_42_-S100A9 amyloids – in blue. *z/m* for Aβ_42_ and S100A9 monomers were calculated by using their amino acid sequences at pH 7.4 and shown by horizontal lines in corresponding colours. The median values for each type of particles are indicated in figure legend.

**Figure S7.**
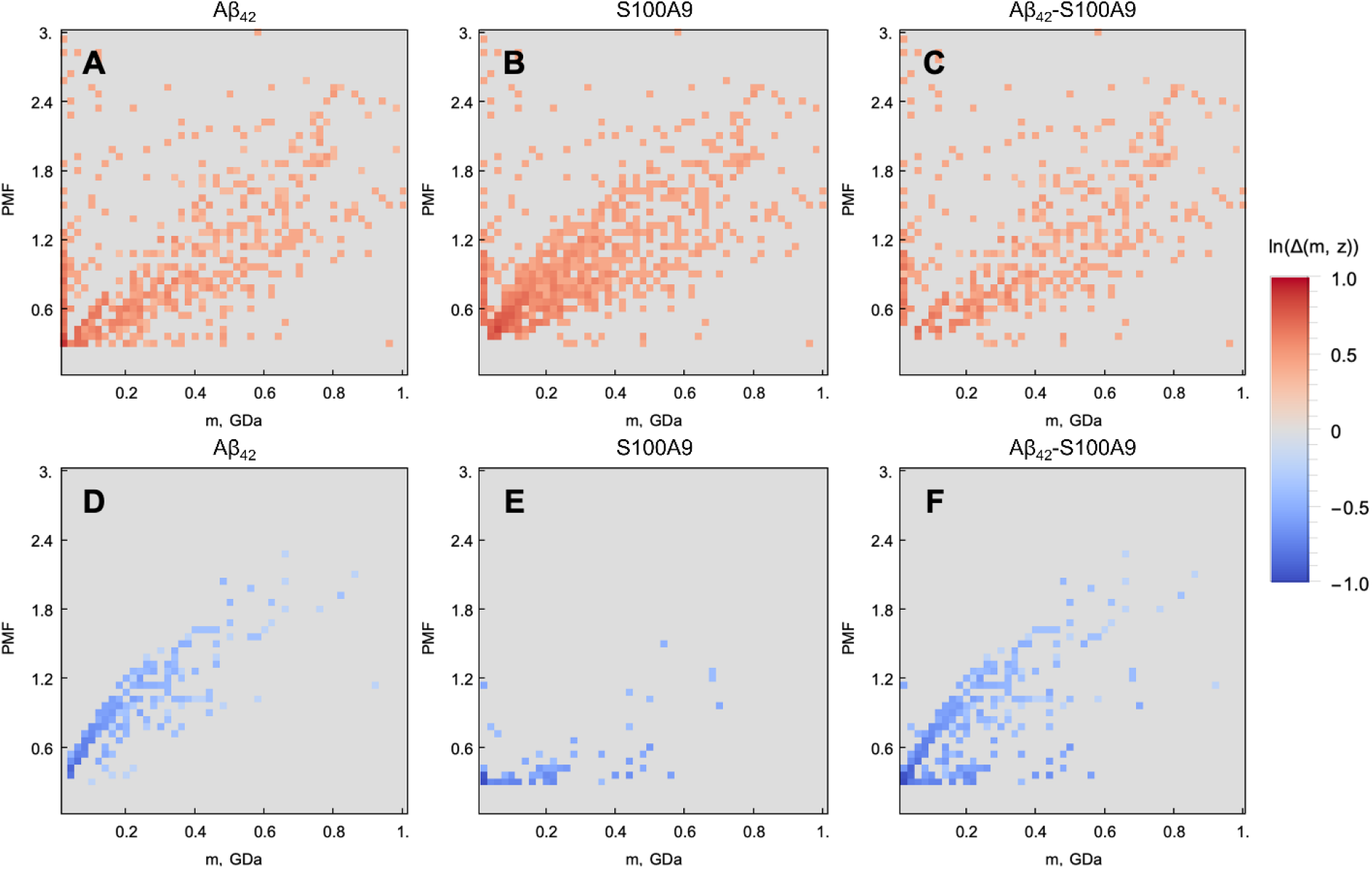
Difference maps derived by comparing CDMS data of Aβ_42_-S100A9, Aβ_42_ and S100A9 amyloid samples. Difference maps showing enriched populations of particles (in red) between the following amyloid samples: (A) Aβ_42_-S100A9 and Aβ_42_, (B) Aβ_42_-S100A9 and S100A9 and (C) Aβ_42_-S100A9 and Aβ_42_ plus S100A9 samples filtered together, see ESI Material and Methods. Difference maps showing depleted populations of particles (in blue) between the following amyloid samples: (D) Aβ_42_-S100A9 and Aβ_42_, (E) Aβ_42_-S100A9 and S100A9 and (F) Aβ_42_-S100A9 and Aβ_42_ plus S100A9 samples filtered together. The colour scales are shown on the right. 30 μM of each polypeptide were incubated individually or in mixture with each other for 24 h in PBS, pH 7.4 and 42°C.

**Figure S8.**
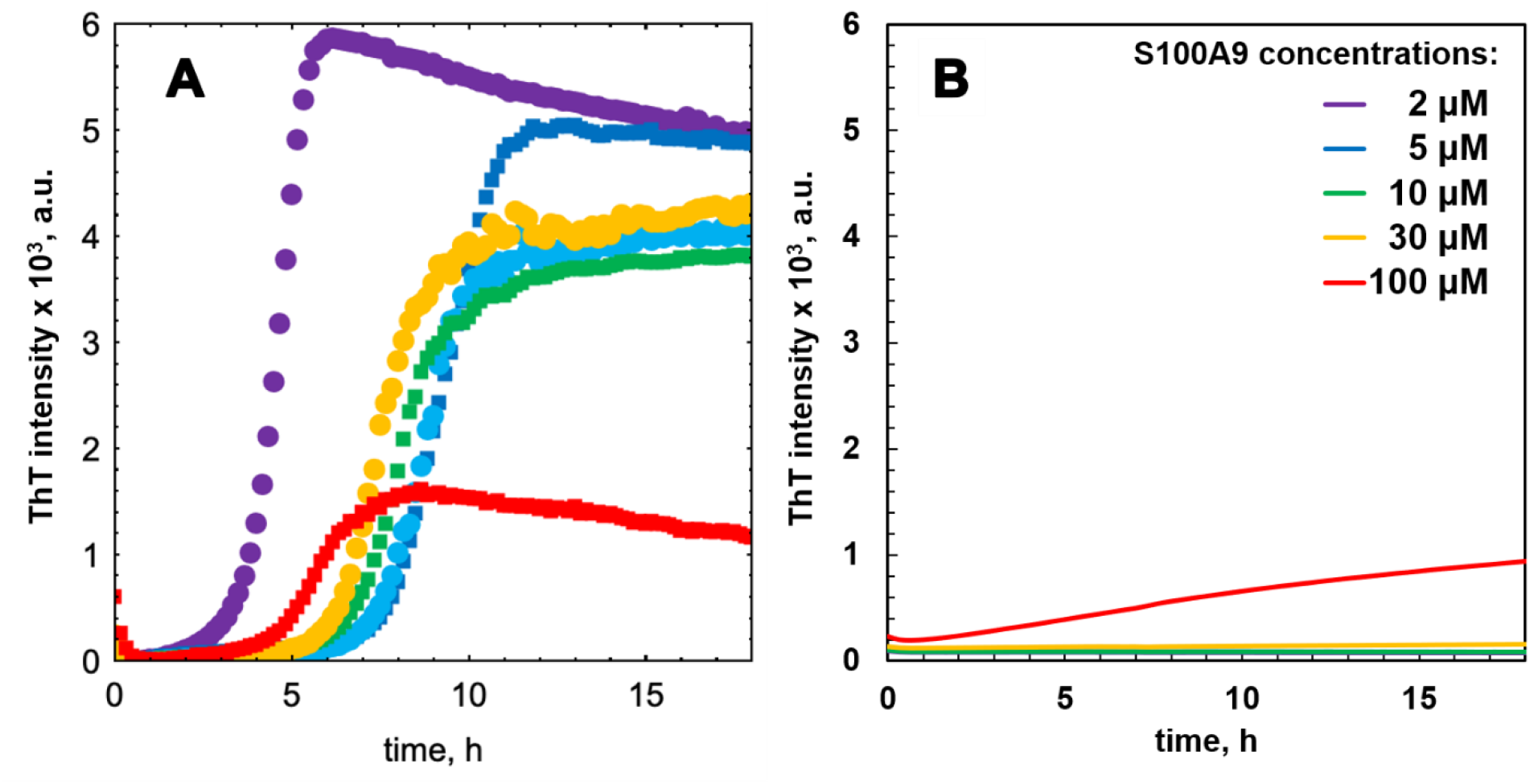
Amyloid fibrillation of Aβ_42_, S100A9 and Aβ_42_-S100A9 mixture monitored by ThT fluorescence assay. (A) Fibrillation kinetics of Aβ_42_ and S100A9 mixtures monitored by ThT fluorescence. (B) Kinetics of S100A9 amyloid formation monitored at different concentrations as indicated in corresponding colour coding in the caption to Fig. 3B. 30 μM Aβ_42_, PBS, pH 7.4 and 42°C.

**Figure S9.**
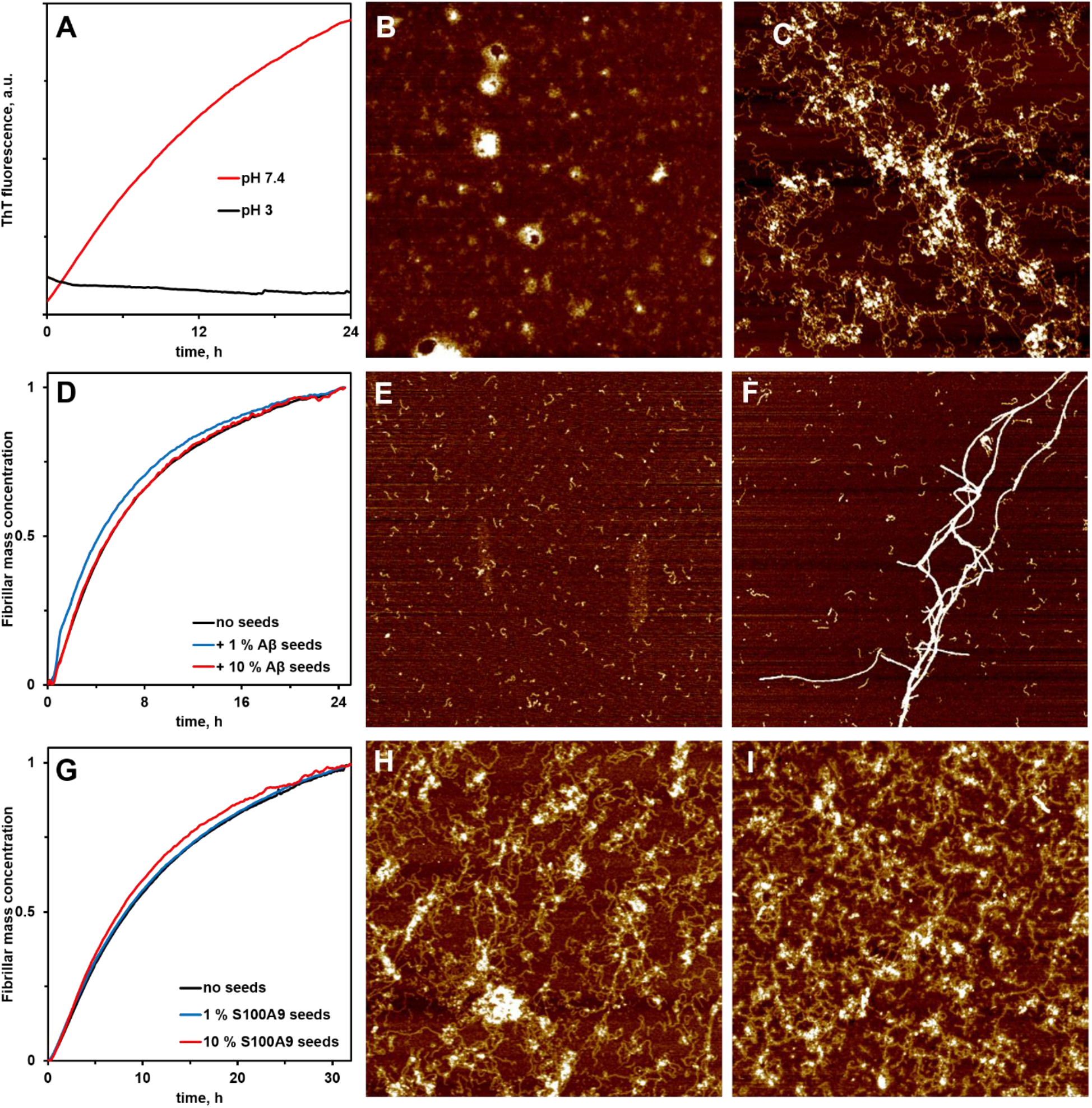
S100A9 does not form amyloids at pH 3 and its amyloids are not seeded by either its own seeds or Aβ_42_ fibril cross-seeds. (A) S100A9 amyloid formation kinetics monitored by ThT fluorescence in 10 mM HCl, pH 3 (black line), PBS, pH 7.4 (red line) and 42°C. AFM images of (B) S100A9 aggregates observed at pH 3 and (C) pH 7.4 after 24 h incubation, 100 μM S100A9. (D) S100A9 amyloid formation kinetics monitored by ThT fluorescence in the absence (black line) and the presence of Aβ_42_ seeds (in corresponding colours indicated in caption). 200 μM S100A9. AFM images of (E) 20 μM S100A9 in the absence and (F) in the presence of 2 μM Aβ_42_ seeds after 8 h incubation. (G) S100A9 amyloid formation kinetics monitored by ThT fluorescence in the absence (black line) and the presence of S100A9 seeds (in corresponding colours indicated in caption). 100 μM S100A9. AFM images of (H) S100A9 in the absence and (I) in the presence of 10 μM S100A9 seeds after 24 h incubation. All experiments were carried out in PBS, pH 7.4 (except pH 3 condition) and 42°C. Scan sizes are 5 x 5 μm.

**Figure S10.**
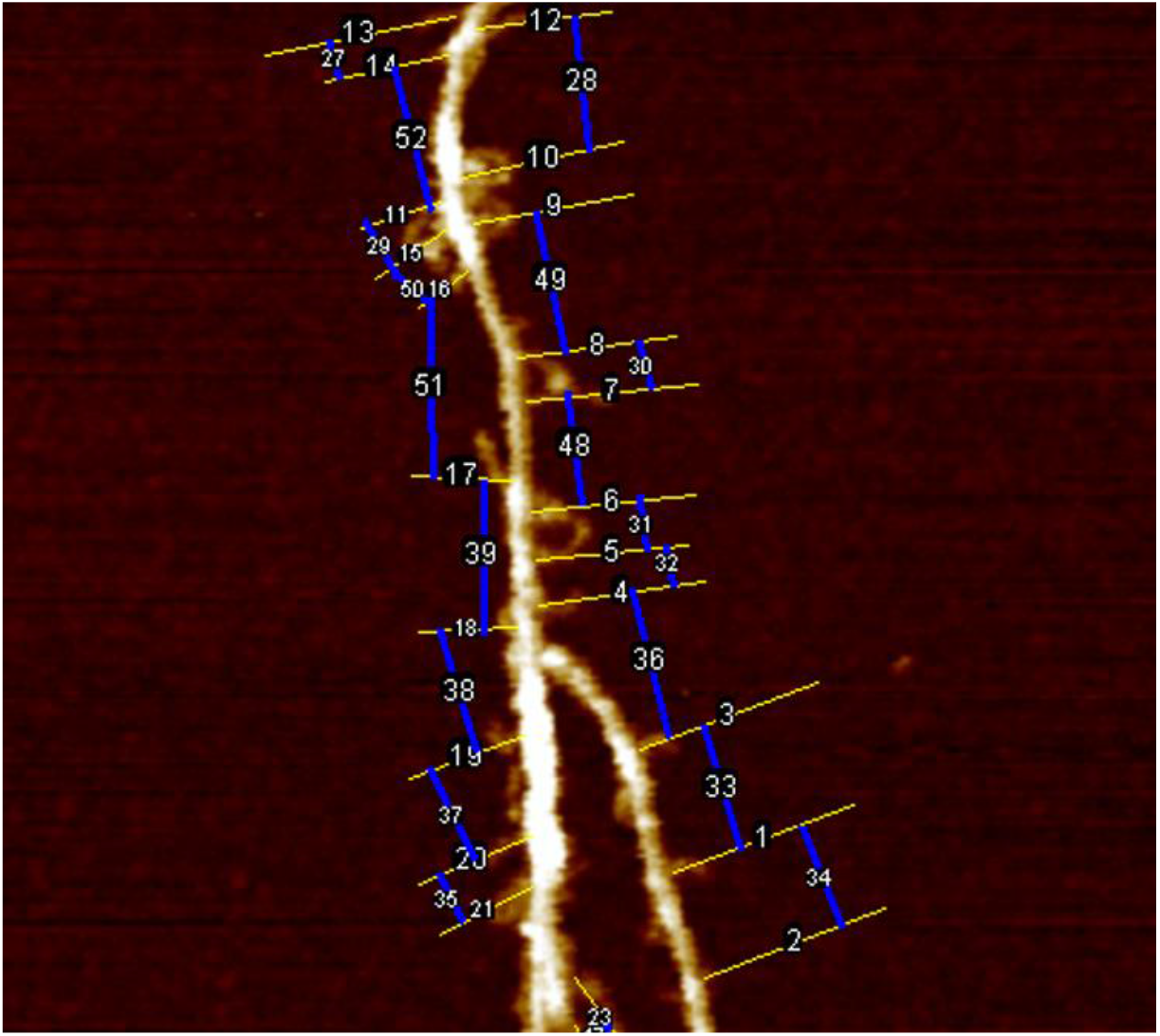
AFM measurements of the distances between S100A9 fibrils templated on the surface of Aβ_42_-S100A9 amyloids. Representative AFM image of Aβ_42_-S100A9 amyloid complex demonstrating the measurements of distances between templated S100A9 filaments. 30 μM of each polypeptide were co-incubated for 24 h in PBS, pH 7.4 and 42°C.

**Table S1.** Statistical analysis of CDMS experimental data.

| | $A\beta_{42}$ | S100A9 | $A\beta_{42}-S100A9$ |
| --- | --- | --- | --- |
| Cumulative probability<sub>m<0.08 GDa</sub> (4 of 50 bins) | 0.37 | 0.45 | 0.43 |
| Cumulative probability<sub>z<0.36 ke</sub> (6 of 50 bins) | 0.1 | 0.55 | 0.2 |

## Notes

### Competing Interest Statement

The authors have declared no competing interest.

